# Versatility of *Campylobacter jejuni* Bf extracellular vesicles in regulating adaptation and virulence under combined thermal and oxidative stress

**DOI:** 10.64898/2026.03.26.714464

**Authors:** Jeanne Malet-Villemagne, Rochelle D’Mello, Yingxi Li, Zoran Minic, Karine Gloux, Florence Dubois-Brissonnet, Bastien Prost, Audrey Solgadi, Christine Péchoux, Vlad Costache, Marianne De Paepe, Zhu Zhang, Gilles Tessier, Jasmina Vidic

## Abstract

The high prevalence of aerotolerant human *Campylobacter jejuni* isolates suggests a correlation between the ability to survive in aerobic conditions, virulence and resistance to harsh stress conditions. However, the mechanisms are still unclear. Here, we investigated the role of bacterial extracellular vesicles (bEVs) in the adaptation of the clinical aerotolerant *C. jejuni* Bf strain to thermal and oxidative stress. We show that *C. jejuni* Bf survives and actively multiplies under this combined stress. Stress exposure induced cell rounding and loss of motility, remodeling of membrane composition, decreased membrane fluidity, and metabolic reprogramming with increased intracellular ATP levels. Lipidomic analyses further revealed that bEVs composition is markedly different from that of the parent membranes indicating that vesicle formation is selective and regulated. Although bEVs were produced in similar amounts under both microaerophilic and stress conditions, stress exposure generated significantly larger vesicles with greater diameter and dry mass, and altered their protein and lipid profiles. bEVs derived from stressed cells showed increased toxicity toward the epithelial barrier of Caco-2 cells. Taken together, these results indicate that *C. jejuni* bEV secretion is part of a survival strategy that connects environmental adaptation with pathogenicity.

**Graphical abstract:** 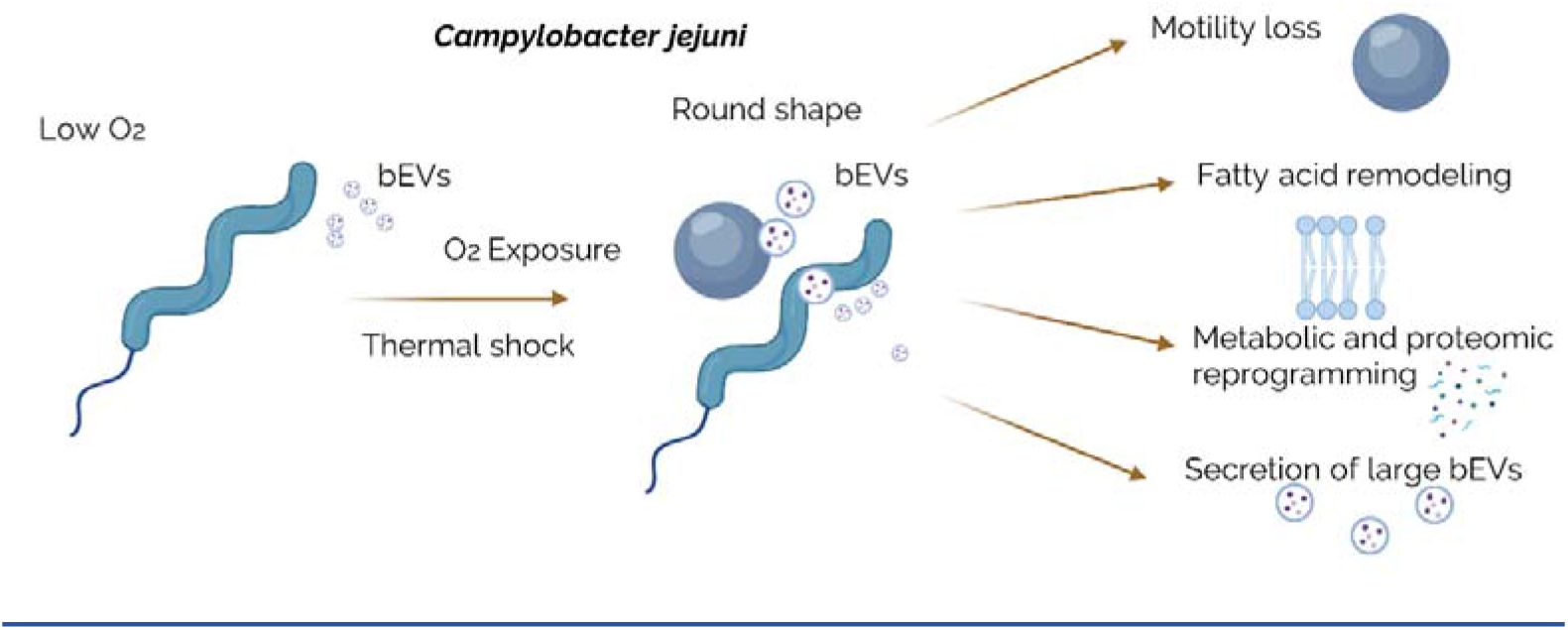

## 1. Introduction

Bacterial infections remain a major challenge in medicine and public health. They are the second-leading cause of death globally, after ischemic heart disease (Murray 2024). Microbes establish infection and survive within their hosts through continual evolutionary adaptation. Additionally, adaptations across diverse biotopes, including animals, food, and water, can enhance bacterial infectivity and enable host switching (Wareth and Neubauer 2021, Kauer, Sapountzis et al. 2025, Zhang, Han et al. 2025). Understanding microbial adaptation to consecutive stress within ecosystems, which allows pathogens to colonize and persist in their hosts, has profound implications for improving public health as highlighted by the WHO’s One Health approach.

*Campylobacter* infection is among the leading causes of foodborne gastroenteritis worldwide. As a commensal organism in the intestinal tracts of food-producing animals, *Campylobacter* readily enters the food chain and is transmitted to humans primarily through the consumption of contaminated food products. Epidemiological studies estimate that 50 to 80% of human campylobacteriosis cases originate from the poultry reservoir (Rasschaert, De Zutter et al. 2020, Authority, Prevention et al. 2024) with *C. jejuni* (~80%) as a leading strain causing human infections *(Hansson, Sandberg et al. 2018)*. Notably, the persistence of *C. jejuni* along the food chain presents a paradox, given its fastidious growth requirements under laboratory conditions and its widespread occurrence in seemingly unfavorable environmental settings (Vizzini, Vidic et al. 2021, Vidic, Auger et al. 2023).

Once chickens become colonized, *C. jejuni* rapidly spreads throughout the flock and persists in the intestinal tract until slaughter (Hermans, Pasmans et al. 2012). During processing, leakage of intestinal material can lead to carcass contamination, with bacterial loads reaching up to 6.5 log CFU/g depending on processing conditions (Rasschaert, De Zutter et al. 2020). In Europe, under the *Campylobacter* process hygiene criterion, official control of poultry samples found that 16.0% exceeded the limit of 3 log CFU/g (Authority, Prevention et al. 2024).

During poultry processing, *C. jejuni* is exposed to various stresses, including heat and cold shocks, acid stress, and oxidative stress. Under acute exposure, the spiral-shaped *C. jejuni* cells become coccoid due to the oxidative damage (Jackson, Davis et al. 2009). This morphological change is one of the events that occur as bacteria enter into a Viable But Non Culturable (VBNC) state, which hinders detection on conventional culture media during quality control procedures (Vizzini, Braidot et al. 2019, Vizzini, Manzano et al. 2021). The phenomenon may explain the continued incidence of campylobacteriosis in developed countries, despite advanced sanitation and food safety standards. At refrigerator temperatures, the membrane fatty acid composition of coccoid cells was shown to be almost identical to that of spiral cells, while intracellular ATP levels can remain high (Hazeleger, Janse et al. 1995). Visualization of VBNC *C. jejuni* cells in aging cell suspensions showed that the outer membrane was lost (Tangwatcharin, Chanthachum et al. 2006). In addition, membrane instability caused by abiotic stress promotes vesicle budding as a way to relieve cell envelope tension or export damaged components (Malet-Villemagne and Vidic 2024). The bacterial production of extracellular vesicles (bEVs) is part of adaptive and survival strategy in hostile environments, including during food processing or within the host (Gill, Catchpole et al. 2019, Malet-Villemagne and Vidic 2024, Calzuola, Malet-Villemagne et al. 2025).

To cope with abiotic stress, *C. jejuni* cells create proteins as a response to new conditions. Specifically, the bacterium produces heat shock proteins (e.g., GroEL, ClpB, DnaK) in response to thermal and acid stress (Stintzi 2003, Kim, Chelliah et al. 2021), and detoxification enzymes (e.g., KatA, SodB, AhpC, Gpx, Ccp) to counter oxidative stress (Baillon, van Vliet et al. 1999, Stintzi 2003, Garenaux, Guillou et al. 2008, Oh, McMullen et al. 2015). In addition, the multifunctional protein, polynucleotide phosphorylase (PNPase), enables the long-term survival of *C. jejuni* at common refrigeration temperatures (Haddad, Burns et al. 2009). While earlier studies often examined only single types of stress (acidic, oxidative, or thermal), recent research has begun to investigate the impact of successive stresses encountered during poultry slaughter and processing (Duqué, Rezé et al. 2021). This approach can better predict *Campylobacter* behavior, as cell history plays an important role in adaptation by introducing physiological responses involved in co-selection mechanisms, including co-resistance, co-regulation, biofilm formation, or entry into a VBNC state (Huo, Xu et al. 2024, Léguillier, Pinamonti et al. 2024). While many stress conditions decrease ATP due to growth inhibition or metabolic slowdown, transient or mild stress conditions can induce short-term increases or stabilization of ATP levels associated with adaptive metabolic responses and subsequent behavior regarding additional stresses (Cohn, Ingmer et al. 2007, Dufour, Stahl et al. 2013, Kim, Chelliah et al. 2021).

Resistance to environmental stresses during processing varies among strains (Robyn, Rasschaert et al. 2015). Not all subtypes survive the abattoir environment and the genetic diversity of *Campylobacter* decreases through carcass chilling during processing (Hunter, Berrang et al. 2009). Importantly, the cold-tolerant *C. jejuni* food isolates usually exhibit elevated aerotolerance (Hur, Kim et al. 2024). The majority of *C. jejuni* clinical isolates and those from retail chicken meat show high aerotolerance and can survive more than 24 hours of aerobic exposure (Oh, Chui et al. 2018, Mouftah, Cobo-Díaz et al. 2021). Therefore, to decrease the prevalence of human Campylobacterioses, it is necessary to understand the mechanism of adaptation to environmental stress in aerotolerant *Campylobacter* strains (Pokhrel, Thames et al. 2022).

Here we show how consecutive stresses, mimicking poultry processing steps, affect the fitness and virulence of *C. jejuni*. We used the clinical Bf strain, previously shown to tolerate aerobic environments (Rodrigues, Pocheron et al. 2015) and to survive starvation in sterile water by entering a non-coccoid VBNC state (Federighi, Tholozan et al. 1998). We focus on structural, lipidomic and proteomic cell adaptation to combined thermal and oxygen stress, as well as the interaction of adapted cells with a human intestinal cell line Caco-2. We further show the role of extracellular vesicles in the adaptive process to abiotic stress employed by *C. jejuni* and its virulence.

## 2. Material and Methods

### 2.1 Bacterial strains and culture conditions

*Campylobacter jejuni* strain Bf (Federighi, Tholozan et al. 1998, Bronnec, Turoňová et al. 2016) was stored at −80°C in brain heart infusion (BHI, Biomérieux, France) supplemented with 20% (v/v) glycerol. The strain was cultured on Columbia agar plates supplemented with 3% defibrinated horse blood for 24-48h at 37°C under microaerobic conditions. Then, a liquid culture was prepared from a single colony of the plate in BHI broth (Biomérieux) supplemented with 50 µmol/L hemin (Sigma-Aldrich) and incubated for 18h at 37°C under microaerobic atmosphere and shaking.

### 2.2 Stress induction protocol

The experimental procedure was conducted as described previously (Duque, Haddad et al. 2019). Briefly, *C. jejuni* Bf was submitted to consecutive stresses inspired from conditions encountered during poultry slaughter. The experimental protocol was thus designed to mimic those consecutive stresses. First, 10 mL of the initial culture was immersed in a hot water bath (50°C) for 5 min and placed in a freezer (−20°C) for 4 min to provoke thermal shock. Then, oxygen was introduced in the culture using a 10 mL syringe and the culture tube was placed at 4°C with the cap open (to pursue oxygen exposure) for different times depending on the experiment. **Fig. 1** illustrates the stress protocol used and the analyses performed in this study.

**Figure 1.**
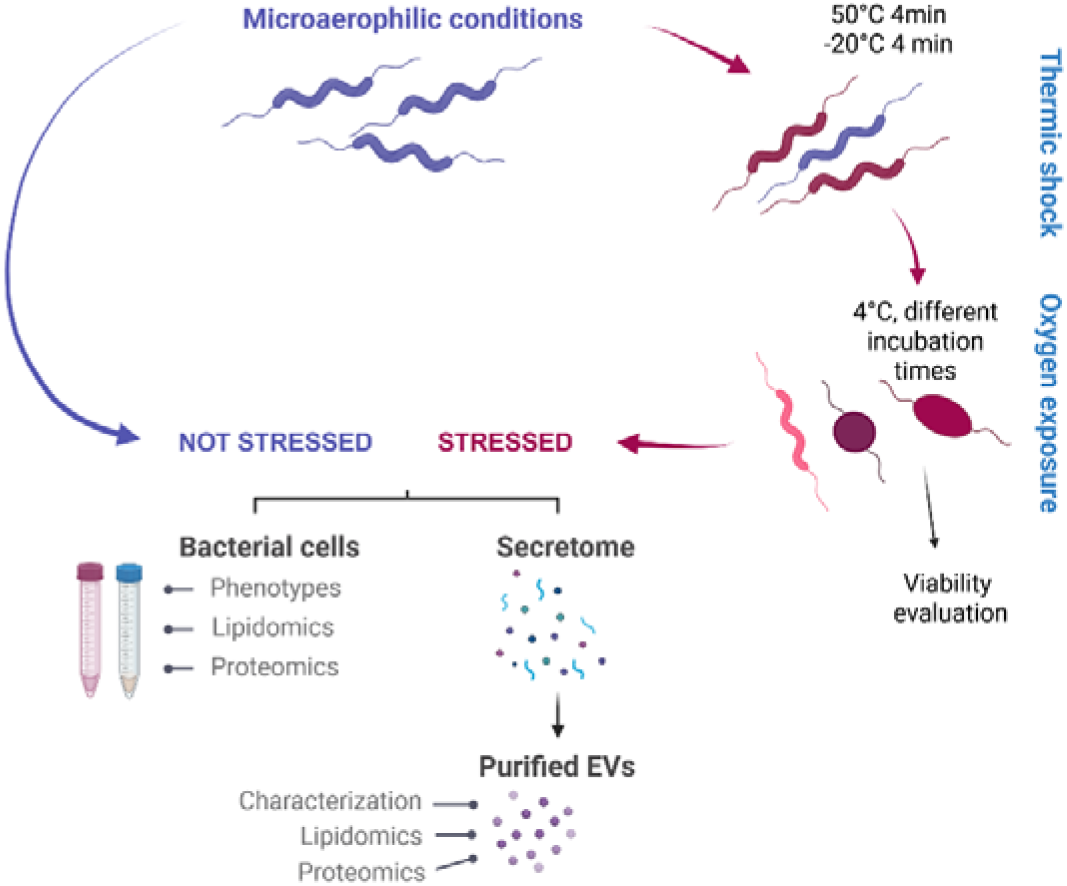
Schematic illustration of the tests performed in this study. The protocol established to mimic consecutive stresses encountered by *C. jejuni* during poultry slaughter included thermal and oxidative stresses. Stressed and unstressed bacterial cells, as well as their secretomes and bEVs, were characterized and analyzed for their lipidomic and proteomic contents.

### 2.3 Optical microscopy for bacteria visualization

Optical microscopy was performed to visualize the bacterial cells exposed to the stress protocol. Observations were made with an Axio-Observer Z1 inverted microscope (Zeiss, Oberkohen, Germany) equipped with a Zeiss AxioCam MRm digital camera, using a x100 oil immersion objective lens. Images were acquired using the ZEN software package (Zeiss, Oberkohen, Germany) and analyzed using the Fiji software ((Schindelin, Arganda-Carreras et al. 2012); http://rsb.info.nih.gov/ij/).

### 2.4 Bacterial viability assay

To assess viability after thermic and oxidative stress, *C. jejuni* Bf was first cultured for 16h at 37°C in microaerophilic conditions and then submitted to the stress protocol described before. The viable population was quantified using the plate enumeration method. Briefly, 10-fold serial dilutions of the bacterial suspensions were made before stress, after thermic shock, and after 4h of air exposure and 100µL of each suspension was spread on BHI supplemented with 1% horse blood. All experiments were made in triplicate, with 2 plates/dilution to allow statistical conclusions.

### 2.5 Intracellular ATP quantification

The intracellular ATP content of bacteria was determined using the luciferase-based BacTiter-Glo Microbial Cell Viability Assay (Promega, #G8230). As per the manufacturer instructions, 100 µL of each culture was mixed in a black, clear-bottomed 96-well plate (Greiner Bio-one, 655090) with 100 µL of room-temperature-equilibrated Bactiter-Glo reagent, and incubated at 25°C in the dark under shaking (160 rpm) for 5 min. Relative luminescence units (RLU) were then recorded in a TECAN Spark® multimode microplate reader (Tecan Life Sciences, Switzerland). Fluorescence values were normalized to an OD_600_ = 1 to allow comparison between samples.

### 2.6 Electrophoretic motility and hydrophobicity

The electrophoretic motility of *C. jejuni* cells and bEVs was analyzed by zeta potential measurements using a Zetasizer Pro (Malvern Panalytical Instrumentation, UK) and the disposable capillary cells DTS1070. At least three measurements were performed on each sample and results are presented as mean ± SD. The hydrophobicity of the *C. jejuni* cells’ surface was assessed using the method of microbial adhesion to solvents by comparing the cell affinity to the polar (PBS) and the non-polar solvent (xylene) as previously described (Shaffique, Imran et al. 2022) with some modifications. Xylene (300µL) was added to the test tubes containing 2mL of washed bacterial cells (~10^8^ CFU/mL) in PBS. After a 10 min incubation at 30°C, upon agitation on vortex for 120 sec, the suspensions were allowed to separate into the polar and non-polar phases. The PBS polar phase was carefully removed with a Pasteur pipette and transferred to an UV-Vis cuvette. Optical density was measured at 400 nm using a Novaspect II spectrophotometer (France). The experiments were performed with three biological replicates.

### 2.7 Permeability test

To assess the permeability of *C. jejuni*’s envelope, strains were grown in BHI supplemented with 50 µmol/L hemin (Sigma-Aldrich) at 37°C overnight. Cultures were pelleted by centrifugation and washed in PBS. The OD_600_ was adjusted to 0.5 with PBS, and 200µL aliquots were distributed to a 96-well cell culture plate. Then, ethidium bromide was added into the wells with a final concentration of 1 µg/mL (Rodrigues, Ramos et al. 2011). Fluorescence was measured using the Tecan Spark multimodal reader (Tecan Life Sciences, France) with the 530 nm band-pass and the 600 nm high-pass filters as the excitation and detection wavelengths, respectively. Fluorescence was measured every 60 sec for 120 min at 37°C. All assays were performed with three biological replicates.

### 2.8 Bacterial EVs isolation and purification

BEVs were collected and purified from non-stressed and stressed *C. jejuni* culture supernatants using a protocol adapted from (Sausset, Krupova et al. 2023). Briefly, bacterial cells were removed by a 5min centrifugation at 5750g and filtered through a 0.22µm membrane filter. The supernatant was then ultra-centrifuged at 120 000 g for 2 h in a Beckman XL-90 ultracentrifuge with a SW41 rotor to pellet down the vesicles. The resulting vesicle pellet was suspended in PBS, and then added on a three-layer iodixanol discontinuous gradient of 45%, 26% and 10% (w/v) in SM Buffer (50 mM Tris pH7.5, 100 mM NaCl, 10 mM MgCl_2_). Tubes were then ultra-centrifuged at 200 000 g for 5h in a Beckman XL-90 ultracentrifuge with a SW55 rotor and purified bEVs were collected and stored at 4°C.

The concentration of bEVs was determined by the rate of Brownian motion in a ZetaView particle tracking analyzer (Excilone Services, France), equipped with fast video capture and particle-tracking software (ZetaViewer, Excilone Services, France). Samples were diluted in PBS before analysis (usually 1:50) and 1 mL of the diluted bEV solution was discharged into the system’s chamber. The size distribution of bEV samples was characterized using dynamic light scattering (DLS) with a Zetasizer Pro (Malvern Panalytical, Malvern, UK). The scattering intensity data was processed using the instrument’s software to obtain the size distribution of the particles. NanoFCM (High Sensitivity Flow Cytometry for Nanoparticle Analysis) was used to assess the purity, concentration, and mean sizes of EV samples. It was performed on a nanoFCM flow nano-analyzer (NanoFCM Co.) following the manufacturer’s instructions to confirm particle size. The instrument was calibrated for particle size with 200 nm polystyrene beads and a silica nanosphere mixture (pre-mixed silica beads with diameters of 68, 91, 113, and 151 nm, currently called “NanoFCM Quality Control Nanospheres”, provided by nanoFCM). 20 µL of each bEV preparation was diluted with sterile PBS in the 1:10 to 1:2000 range allowing the detection of at least 500 particles per acquisition. Events were recorded twice for each biological sample during 60 sec. The flow rate and side scattering intensities were converted into the corresponding particle sizes using the nanoFCM software. MemGlow™ 488 (Cytoskeleton, USA) labelling was used to assess bEV purity by labelling membranes. Live/dead Bacterial Viability kit was used to observe the presence of internal (Syto9) and external (Propidium Iodide) DNA carried by the bEVs.

### 2.9 Vesicle imaging and dry mass measurement using QLSI

The dry mass measurements were carried out using a custom-made quantitative phase imaging system, under the assumption that optical phase distortions can be considered proportional to the dry mass, i.e., the mass of the solute without water (Bon, Maucort et al. 2009, Gentner, Rogez et al. 2024). This Quadriwave Lateral Shearing Interferometry (QLSI) setup, based on an inverted microscope (Olympus IX-71) has been described in (Gentner, Rogez et al. 2024). A visible supercontinuum laser (SMHP-Visible, Leukos) combined with a bandpass filter (FBH450-40, Thorlabs) was used to isolate a Δλ = 40 nm wavelength band centered at λ = 450 nm to illuminate the samples. The source was coupled into a 20 m-long multimode optical fiber (FG105LVA, Thorlabs) and collimated by an aspheric lens (f’ = 4.34 mm, Thorlabs) to illuminate the sample with a plane wave of high spatial coherence (N.A. = 0.04). The light was collected by a 60×, N.A. = 1.4 oil immersion objective (Olympus PlanApo). The back focal plane of this objective was imaged using a f’ = 100 mm Achromatic Doublet (Thorlabs), and a disc-shaped gold filter (100 µm diameter, Au thickness100 nm) was placed in the Fourier plane to attenuate the central order and increase phase sensitivity whenever necessary (see (Gentner, Rogez et al. 2024)). The samples were then imaged on a custom-made high resolution wavefront sensor (WFS) based on QLSI (Bon, Maucort et al. 2009). A two-dimensional 20 μm-step 0−π phase-only checkerboard grating QLSI mask (PhiMask, Idylle), was reimaged a few millimeters (d = 1.6 mm) before the camera. In order to reduce system vibrations, we used a passively cooled (i.e., no fan) ZWO ASI174MM camera.

The setup was used without the dot-shape spatial filters in the Fourier plane, because the standard QLSI was sensitive enough to measure the mass of the vesicles. The ZWO ASI174MM camera, which has no fan in the body, was used to avoid the fan-induced vibration in the system. A visible supercontinuum laser (SMHP-Visible, Leukos) combined with a bandpass filter (FBH450-40, Thorlabs) was used to isolate a Δλ = 40 nm wavelength band centered at λ = 450 nm. Both sources are coupled into a 20 m-long multimode optical fiber (FG105LVA, Thorlabs) and collimated by an aspheric lens (f’ = 4.34 mm, Thorlabs) to illuminate the sample with a plane wave with high spatial coherence (N.A. = 0.04).

The light was then collected by a 60×, N.A. = 1.4 oil immersion objective (Olympus PlanApo). The pupil plane of this objective is not directly accessible as it lies inside it and is reimaged outside the microscope using a f’ = 100 mm Achromatic Doublet (Thorlabs). After this, the samples were imaged on a custom-made high resolution wavefront sensor (WFS) based on QLSI (Berto, Rigneault et al. 2017, Brasiliense, Audibert et al. 2022). This WFS consists in a two-dimensional 20 μm-step 0−π phase-only checkerboard grating (PhiMask, Idylle), reimaged a few millimeters (d = 1.6 mm) before the sCMOS camera (ZWO ASI174MM).

### 2.10 Imaging of bacterial cells and bEVs by Transmission Electron Microscopy (TEM)

Samples of bacteria were fixed with 2% glutaraldehyde in 0.1 M Na cacodylate buffer pH 7.2, and contrasted with Oolong Tea Extract (OTE) 0.2% in cacodylate buffer, postfixed with 1% osmium tetroxide containing 1.5% potassium cyanoferrate, gradually dehydrated in ethanol (30% to 100%) and substituted gradually in a mix of ethanol-epon and embedded in Epon (Delta microscopie, France). Thin sections (70 nm) were collected onto 200 mesh copper grids, and conterstained with lead citrate. Grids were examined with the Hitachi HT7700 electron microscope operated at 80kV (Milexia, France), and images were acquired with a charge-coupled device camera (AMT).

For the TEM imaging of bEVs, they were also fixed with 2% glutaraldehyde in 0.1 M Na cacodylate buffer pH 7.2 and then directly adsorbed onto a carbon film membrane on a 300-mesh copper grid, stained with 1% uranyl acetate, dissolved in distilled water, and dried at room temperature. Grids were examined with the Hitachi HT7700 electron microscope operated at 80kV (Milexia, France), and images were acquired with a charge-coupled device camera (AMT).

### 2.11 Imaging of bacterial cells by Scanning Electron Microscopy (SEM)

After overnight incubation, 2 mL of culture was gently pelleted (2000 g, 4 min) to 50 µL that was fixed in 2 mL 2 % glutaraldehyde buffered with sodium cacodylate 0.2 M, during 2 h at room temperature then overnight at 4°C, in a 24 multi-well plate (TPP 92024, Switzerland) that contains 1 cm x 1 cm glass slides at the bottom of the well. Samples attached to the glass slides were rinsed twice for 5 min in 0.2 M sodium cacodylate buffer, then in successive baths of ethanol (50, 70, 90, 100, and anhydrous 100 %), and finally dried using a Leica EM300 critical point apparatus with slow 20 exchange cycles, with a 2 min delay. Samples were mounted on aluminum stubs with adhesive carbon (EMS, LFG France) and coated with 6 nm of Au/Pd using a Quorum SC7620, 50 Pa of Ar, 180 s of sputtering at 3.5 mA. Samples were observed using the SE detector of a FEG SEM Hitachi SU5000, 2 KeV, 30 spot size, 5 mm working distance, located on the MIMA2 core facility, INRAE, Jouy-en-Josas, France; https://doi.org/10.15454/1.5572348210007727E12.

### 2.12 Proteomic studies

For proteome analysis, the samples were concentrated 6.5 times on 10kDa filters to allow for protein quantification by the Bradford assay using a protein assay kit (Thermofisher Scientific) following the manufacturer’s protocol. The assay absorbance was measured with a spectrophotometer (Novaspec III, Biochrom) at 595 nm. Then, aliquots of 40µg of protein was further processed with a modified filter-aided sample preparation (FASP) protocol for proteomic analysis (Wisniewski, Zougman et al. 2009). Briefly, bEVs samples were diluted and incubated for 15min in 200µL of lysis buffer (8M urea, 25 mM HEPES, pH = 8.0, 5% glycerol, 1mM DTT, 0.2% DDM, 1:200 v/v protease inhibitor) before being transferred to a 10kDa molecular weight cut-off (MWCO) filter (MRCPRT010, Millipore). Sample volume was reduced to 50µL prior to the addition of 100µL denaturation buffer (8M urea, 25mM HEPES, pH = 8.0). Again, the sample volume was reduced to 20µL by centrifugation at 14,000g. After that, proteins were reduced by adding 4mM Tris(2-carboxyethyl)phosphine (TCEP) (Thermo Fisher Scientific) in denaturation buffer and incubation for 30 min at RT followed by centrifugation for 30 min at 14,000 g. Proteins were then alkylated using a 40min RT incubation with 20mM iodoacetamide (IAA) in 100µL denaturation buffer followed by a 30-minute centrifugation at 14,000 g. Next, digestion buffer was added (0.6% v/v glycerol, 25 mM HEPES, pH = 8.0), and the remaining urea was washed by a centrifugation step before the addition of MS-grade trypsin/Lys-C mix (Promega) with a 1:150 enzyme to protein ratio. The proteolytic digestion was performed in the dark under agitation (600 rpm) at 37°C for 12 hours. Peptides were eluted as filtrate by centrifugation at 14,000 g for 30 min, and then acidified with 2% (v/v) formic acid and desalted for MS analysis using Pierce C-18 spin columns (Thermofisher Scientific). Purified peptides were vacuum-dried (Savant SPD111V SpeedVac Concentrator, Thermo Scientific) and reconstituted with 20µL of MS grade water with 0.1% formic acid for MS analysis.

Proteomic analysis was carried out as described previously (Li et al., 2024, DOI: 10.1038/s41598-024-65417-2). Approximately 4⍰µg of protein content, quantified using the Bradford assay, were injected in an UltiMate 3000 nanoRSLC (Dionex, Thermo Fisher Scientific, Mississauga, ON, Canada) couplet to an Orbitrap Fusion mass spectrometer (Thermo Fisher Scientific, Mississauga, ON, Canada) for proteomic analysis. The mass spectrometry data were processed for protein identification and quantification using MaxQuant 2.6.4.0 (Cox et al., 2011, DOI: https://doi.org/10.1016/j.ecoenv.2026.119859). Acquired peptide spectra were matched against the *Campylobacter jejuni* protein database from NCBI (taxonomy ID: 197), 1737 reviewed and 78037 unreviewed entries as of 10.10.2024.

A quantitative bioinformatics analysis of *C. Jejuni* whole cell samples was conducted using R programming language in RStudio (2024.04.v2). To visualize common and unique proteins, Venn diagrams were created using the ggVenn package in R where unique proteins were defined as proteins that were present in at least 2/4 samples of a particular group. To comparatively visualize the whole proteome, a principal component analysis was conducted. Missing LFQ intensity data was first imputed. The data was then log2 normalized and scaled. The components were then calculated using the prcomp() function from the stats package. Principal component 1(PC1) and principal component 2(PC2) were plotted using the ggplot2 package.

Additionally, a comparison of the log_2_ normalized data to common proteins was performed by conducting a student’s t test with the t.test() function from the stats package. The log2 fold change was also calculated. Both p-value and fold change parameters, stressed and non-stressed common proteins from *C. Jejuni* whole cells, were plotted to create a volcano plot. Overall protein significance was defined as having a p value of <0.05 and a minimum fold change (stressed/non-stressed) of 1.5.

*C. jejuni* bEV samples were also analyzed using RStudio. Data was imported and filtered as described above. A Venn diagram was also created for bEV using the ggVenn package as described above. Kegg Pathway enrichment analysis was also conducted using the proteins identified in stressed and non-stressed conditions using Eggnog mapper 2.1.12. First, IDs were mapped using the uniport ID mapping tool (https://www.uniprot.org/id-mapping). Results were downloaded and compressed into a fasta file and uploaded into EggNog Mapper (http://eggnog-mapper.embl.de/) for Kegg pathway analysis.

### 2.13 Lipidomic study

To obtain lipid fractions, bacterial cells or purified vesicles were freeze-dried and extracted with 4.75 ml of chloroform-methanol 0.3% NaCl (1:2:0.8 v/v/v) at 80°C for 15 minutes. Solutions were vortexed for 1 hour at room temperature and then centrifuged at 4,000rpm for 15 minutes at room temperature. Supernatants were collected, and the debris was re-extracted with 9.5 ml of the same mixture with 30 seconds of vortexing. After the second centrifugation, supernatants were pooled and diluted with 2.5 ml each of chloroform and 0.3% NaCl solution. Phase separation was achieved by centrifugation at 4,000rpm for 15 minutes at room temperature. The upper phase was discarded, and the chloroform phase was evaporated to dryness under a nitrogen stream and stored at −20°C. Phospholipid analysis was performed on Dionex Ultimate 3000 RSLC system (Thermo Fisher Scientific) equipped with two quaternary pumps, an autosampler, and a column oven. The RSLC system was coupled online to an LTQ Orbitrap Velos Pro mass spectrometer equipped with a linear ion trap and an orbital trap analyzer (Thermo Fisher Scientific, Waltham, MA, USA). Lipid separation was performed as previously described (Moulin, Solgadi et al. 2015). Prior to injection into the chromatographic system, the dry pellets were solubilized in appropriate volumes of chloroform to achieve comparable concentrations (60 mg/mL for bacterial samples) and in a minimal volume of 50 µL for bEVs (6mg/mL-10 mg/mL bEVs). Phospholipids were separated according to their polarity: the retention time provides information on polar head group which is compared to commercial standards. Individual species were then identified under each chromatographic peak. Identification of phospholipids was performed in negative electrospray ionization mode (ESI-) using high-resolution mass detection in full-scan MS1 mode on the Orbitrap. The exact mass obtained in MS1 is used to confirm the class of the phospholipid and to get the number of carbon and the amount of unsaturation of each m/z. Data dependent MS2 fragmentation was also conducted in the linear ion trap for the composition of the fatty acyl chains. MSn fragmentation is not always available due to the low abundance of some ions. In these cases, phospholipids are mentioned as PL(X:n) where PL is the class of the phospholipid, X represents the total number of carbon atoms in the fatty acyl chains and n the total number of unsaturation. Otherwise, phospholipids are reported with their specific fatty acyl chain composition, e.g. PL(16:0-18:1). The sn-1 and sn-2 positions of the acyl chains were not determined.

### 2.14 Fatty acid (FA) identification

FAs profiles were performed using two methods. The first was described previously (Morvan, Halpern et al. 2016). Briefly, bacteria were harvested by centrifugation (5000 g, 5 min) and purified bEVs were freeze-dried before storage at −80°C until further processing. Extraction and trans-esterification of FA were performed directly on bacterial pellets or lyophilized bEVs. Gas chromatography separation was performed by injection of the FA methyl esters in a split-splitless mode of an AutoSystemXL gas chromatograph (PerkinElmer) equipped with a ZB-Wax capillary column (30 m x 0.25 mm x 0,25 mm; Photomenex) and a flame ionization deterctor. FAs were identified based on their retention times. Data were analyzed on TotalChrom Workstation (PerkinElmer). Results are presented as representative gas chromatograph profiles and as FAs peak areas (expressed as a percentage of the total areas of detected peaks).

The second method was used because it allows the extraction of all FA including hydroxylated FA (Guérin-Méchin, Dubois-Brissonnet et al. 1999, Méchin, Dubois-Brissonnet et al. 1999). Briefly, whole-cell FA were first saponified and esterified using methanolic NaOH (1⍰mL of 3.75⍰M NaOH in 50% (v/v) methanol and incubation for 30⍰min at 100⍰°C), followed by treatment with methanolic HCl (addition of 2⍰mL of 3.25⍰M HCl in 45% (v/v) methanol solution and incubation for 10⍰min at 80⍰°C). Fatty acid methyl esters (FAME) were then extracted using a 1:1 (v/v) diethyl ether/cyclohexane mixture. The organic phase was washed with a dilute base solution (0.3⍰M NaOH). Analytical gas chromatography of FAME was carried out in a GC-MS Trace ⍰300 / ISQ 7000 system (Thermo Fisher Scientific) equipped with a BPX70 capillary column (25⍰m, 0.22-mm internal diameter; SGE, Victoria, Australia). The column temperature was initially maintained at 100 °C for 1 min, then gradually increased to 170⍰°C at a rate of 2⍰°C per minute. FA species were identified based on mass spectrometry (MS) databases (Replib, Mainlib, FAME2011) (Touche, Hamchaoui et al. 2023). The relative abundance of each FA was calculated as a percentage of the total FAME peak area. Identified FA species were grouped into the following categories: saturated FA (SFA), unsaturated FA (UFA), and cyclopropane FA (cycloFA).

### 2.15 Membrane fluidity test

The generalized polarization of the lipophilic dye laurdan (6-dodecanoyl-2-dimethylaminonaphtalene), when bound to *C. jejuni* plasma membrane, was used as a measure of bacterial fluidity (Scheinpflug, Krylova et al. 2017). Bacteria were sampled (1mL), collected by centrifugation (5000 g, 5 min), resuspended in PBS and incubated for 10 min in the dark with 10 µM of laurdan (Sigma-Aldrich, #40227) from a 1 mM stock in dimethylformamide (DMF). Then, unbound laurdan was removed using four cycles of centrifugation (8000 g, 5 min) and resuspension in pre-warmed (30°C) 1% (v/v) DMF in mineral water (by vortexing). After final resuspension, technical replicates (200µL) were added to a pre-warmed (30°C) clear-bottom black 96-well plate and laurdan fluorescence was induced at 350 nm and recorded at 450 and 500 nm in a Tecan Spark (Tecan Life Sciences) microplate reader. The general polarization (GP) of laurdan was determined by the formula: GP = (/_450_⍰− ⍰/_500_)/(/_450_⍰+⍰/_500_), where / corresponds to the fluorescence intensity value at the recorded emission wavelength (Scheinpflug, Krylova et al. 2017). An increase of laurdan GP values corresponds to a reduction of membrane fluidity (rigidification).

### 2.16 Caco-2 cell culture and cytotoxicity evaluation

The human colorectal carcinoma cell line (Caco-2, ATCC HTB-37) was cultured following the ATCC recommendation for this cell line, with minor modifications. Caco-2 cells were routinely cultured in DMEM Glutamax growth medium (Gibco, Thermo Fisher Scientific, USA) supplemented with 20% fetal calf serum (FCS, Gibco France), 1% Penicillin-Streptomycin (Sigma-Aldrich, USA) in T25 cell culture flasks at 37°C in a 5% CO_2_ humidified atmosphere. Cells were passaged every 2-3 days. For use in the assays, Caco-2 cells were seeded in 96-well culture plates at a density of 3×10^4^/well and grown until confluence before further treatment.

The cytotoxic effect of BEVs and culture supernatants from *C. jejuni* Bf on the viability of cultured Caco-2 cells was determined by MTT assay. For this, Caco-2 cells were seeded in 96-well plates (Stardedt) and grown 48 hours to confluence before the addition of *C. jejuni* supernatants and bEVs (MOI 20). The blank contained only medium while the positive control contained cells and medium. Supernatants were added at a final concentration of 20% in complete culture medium and the plate was incubated 24 hours at 37°C in a 5% CO_2_ atmosphere. After incubation, the medium was replaced with 100µL media containing MTT reagent (1 mg/mL) and incubated at 37°C in a 5% CO_2_ atmosphere for 1 hour. Subsequently, the media was removed, the plate was dried for 15 min and 100µL of DMSO was added to dissolve the formazan crystals formed. The absorbance of the plate was measured using a microplate reader (TECAN Spark) at 670 nm (for background measurement) and at 560 nm. Cell viability was calculated by removing the blank value to all samples and normalizing treated cells to untreated cells (representing 100% cell viability).

The trans-epithelial electrical resistance (TEER) values reflect the integrity of a given monolayer and are also evidentiary of the polarization of epithelial cells. The TEER across the monolayer of Caco-2 cells was measured first to determine confluency and then to evaluate the cytotoxic effect of *C. jejuni* bacteria and bEVs on intestinal epithelial cells. For the TEER experiment, Caco-2 cells were seeded in 24-well Transwell plates (Corning Costar, 8µm pore size, PET membrane, SURF) and grown 5 days to confluence before the addition of *C. jejuni* cells and bEVs. TEER values between 150 and 600 Ωcm^2^ were considered as indicative of a confluent monolayer, and experimental values observed ranged between those values.

### 2.17 Statistics

Statistical analyses were done using GraphPad Prism 10.2.1 (GraphPad Software). Data obtained from experiments performed with three independent biological replicates (n⍰=⍰3) are represented as mean⍰±⍰standard deviation or median + interquartile range. *p* values ≤0.05 were considered as significant for all analysis performed and were indicated with asterisks: \**p*≤0.05, \*\**p*≤0.01 and \*\*\**p*≤0.001. Additional test details can be found in the *Source Data* file.

## 3. Results

### 3.1 Viability of *C. jejuni* Bf under combined thermic and oxygen stresses

*C. jejuni* encounters several stresses during the poultry slaughter process, particularly during the scalding and chilling steps, which involve heat and cold stress. Afterward, chicken cuts are conditioned and stored under a modified atmosphere, which can cause a significant oxidative stress. We therefore examined the impact of these consecutive stresses on the viability and morphology of the aerotolerant *C. jejuni* Bf (Federighi, Tholozan et al. 1998, Rodrigues, Pocheron et al. 2015). Assuming that *Campylobacter* is not under starvation conditions when attached to chicken carcasses, the tests were performed in BHI broth.

Phase-contrast images of *C. jejuni* suspensions were recorded over one week to track morphological changes in the stressed cells. The initial C. jejuni population (Day 0, Fig. 2A) consisted of typical curved or spiral-shaped cells that moved rapidly in helicoidal or corkscrew patterns through the liquid (**Video S1**). From the first day of incubation under aerobic conditions, coccoid forms appeared and increased in abundance over time at the expense of the spiral-shaped subpopulation (**Fig. 2A**). In addition, bacterial motility progressively decreased, and the cells became immobile, which facilitated microscopic observation (**Video S1, S2**). To confirm the morphological changes, we recorded SEM and TEM images of stressed *C. jejuni*. Untreated Bf cells displayed a one-spiral turn form, as previously observed. (Federighi, Tholozan et al. 1998). After experiencing consecutive stresses, coccoid-like cells were rapidly observed, as shown in **Fig. 2B**. This suggests that morphological adaptations occur as early-stage protective mechanisms against stress. Noteworthy, vacuoles were sometimes found in coccoid cells (TEM images of cellular cross-sections in **Fig. 2B**) further confirming the adaptation strategy. Previously, vacuolated coccoid cells were found in the *C. jejuni* VBNC state, probably assisting the bacterium in compartmentalizing damage and shifting to a dormant morphology (Ng, Sherburne et al. 1985). Quantitative analysis of the population morphology confirmed this progressive shape transformation, as cell length decreased (**Fig. 2C**) while width increased (**Fig. 2D**) over time after treatment.

**Figure 2.**
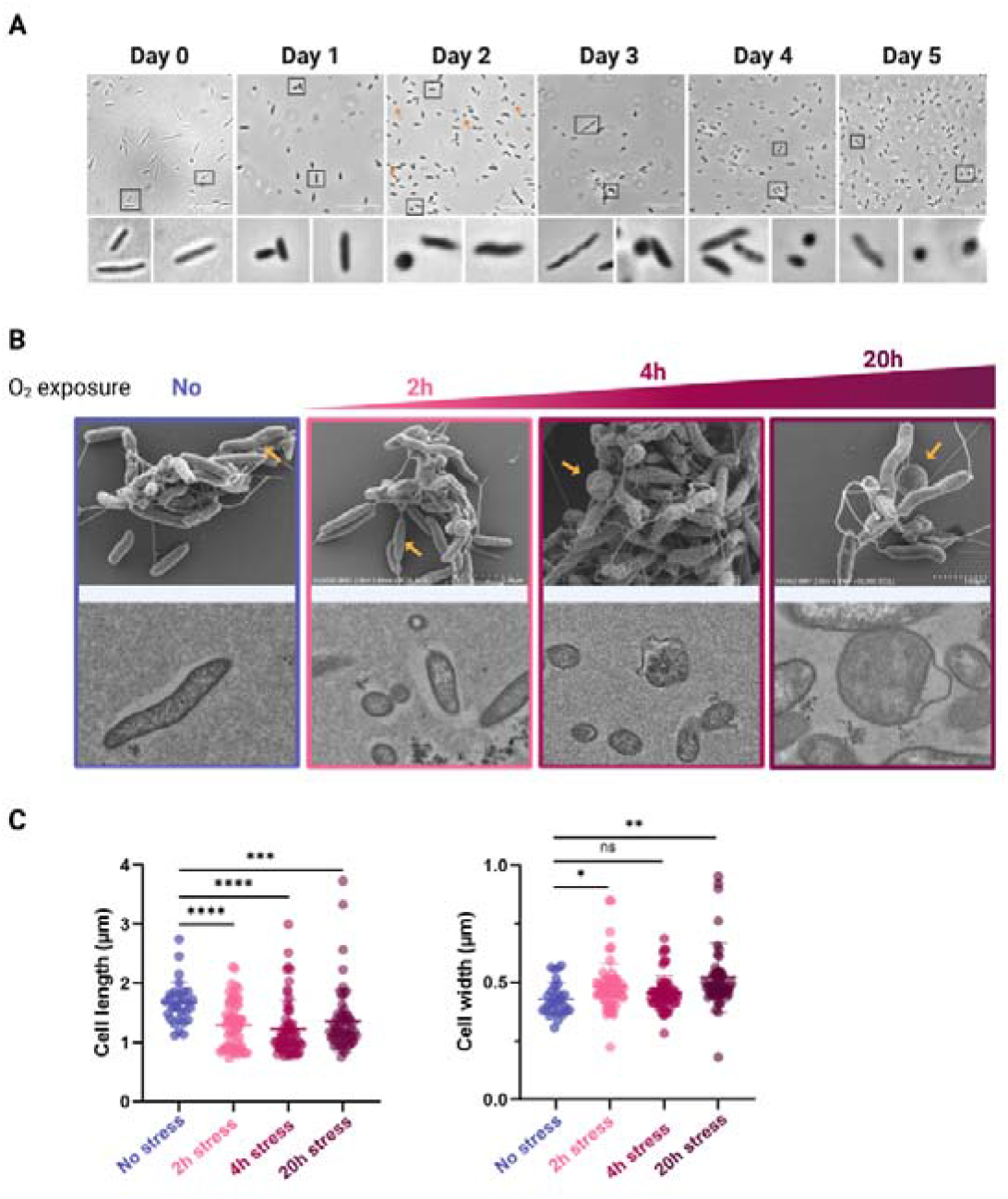
*C. jejuni* Bf survives combined thermal and oxidative stresses by changing its shape from spiral to coccoid. **(A)** Phase-contrast micrographs of *C. jejuni* in BHI over 8 days. Bacteria highlighted in black squares are shown enlarged in the bottom panels. **(B)** Scanning electron microscopy (SEM) and transmission electron microscopy (TEM) images of *C. jejuni* Bf during the 20 hours following stress induction. **(C, D)** Cell length and roundness of *C. jejuni* Bf in BHI. Data are presented as mean ± standard deviation from independent experiments (n=3). Statistical significance was determined by Student’s t test; ns, not significant, * *p* < 0.05, ** *p* < 0.01, *** *p* < 0.001, **** *p* < 0.0001.

We then focused on the potential growth transition of the *C. jejuni* Bf strain from microaerophilic to stress conditions. Bacterial cells grew exponentially under microaerophilic conditions, and at time zero (T_0_) cultures were subjected to successive heat and cold thermal shock, which caused a slight but significant reduction (1 log) in bacterial load, indicating that cell growth was immediately arrested. Surprisingly, although exposed to oxygen, the stressed cells resumed growth (**Figs. 3A** and **3B**). To confirm that stress conditions mimicking poultry slaughtering steps do not induce a dormant state in *C. jejuni* Bf cells, ATP levels were measured in untreated and stressed cells at 4h post-stress. ATP can be detected only in live cells, as it is quickly depleted upon cell death (Eydal and Pedersen 2007), while dormant cells show a reduction in intracellular ATP levels (Ayrapetyan, Williams et al. 2018). The measured ATP levels were higher in stressed cells compared to untreated cells (p<0.05, **Fig. 3C**). Although the role of ATP in regulating stress responses has not been specifically investigated, high ATP level have been shown to enable recovery of dormant cells. We may thus hypothesize that combined thermal and oxygen stress induces adaptations that enable proliferation of Bf cells.

**Figure 3.**
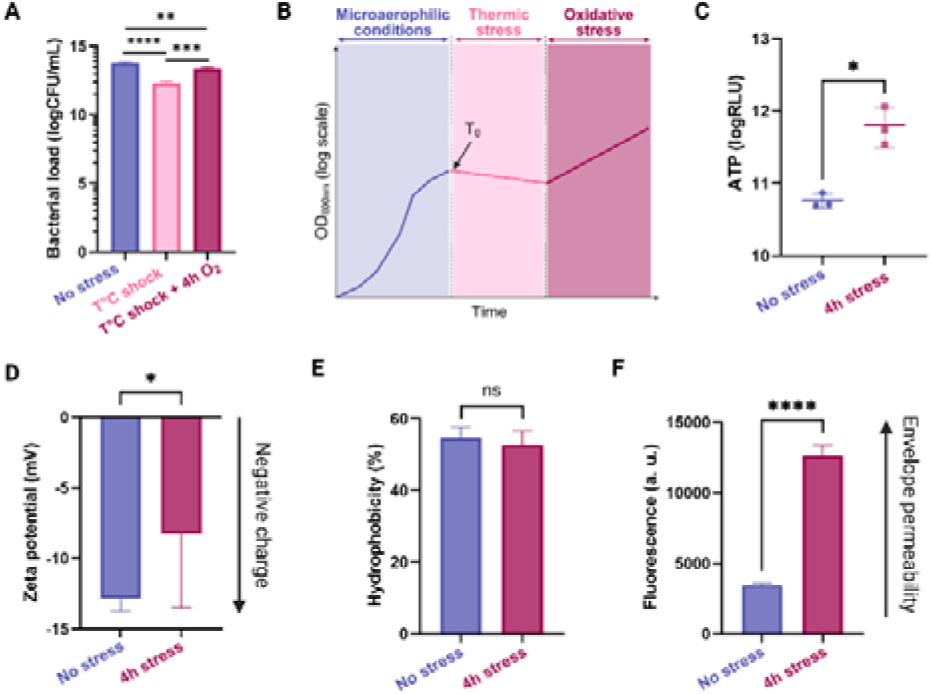
*C. jejuni* Bf multiplies under combined thermal and oxidative stress. **(A)** Bacterial load of *C. jejuni* in CFU/mL enumerated on plates before stress, after thermal shock, and after combined thermal and oxidative stresses. **(B)** Schematic illustration of a growth curve for *C. jejuni* Bf surviving combined thermal and oxidative stress. **(C)** ATP content of *C. jejuni* non-stressed and stressed cells, quantified using a luciferase-based assay. **(D)** Zeta-potential measurements of electrophoretic mobility of bacteria reflecting their surface charge. **(E)** Cell surface hydrophobicity based on the affinity of cells for xylene solvent. **(F)** Bacterial cell wall permeability measured with the ethidium bromide influx assay. Data are presented as mean ± standard deviation from independent experiments (n=3). Statistical significance was determined by Student’s t test; ns means not-significant, * *p* < 0.05, ** *p* < 0.01, *** *p* < 0.001, **** *p* < 0.0001.

To determinate whether the physicochemical characteristics of the cell envelope changed during *Campylobacter* morphological adaptation to stress, we examined the bacterial cell surface. Stressed cells exhibited a less negative surface charge than untreated cells, as determined by zeta potential measurements (−12.9 ± 0.9 mV in control vs −8.3 ± 5.2 mV in stressed cells, **Fig. 3D**), while no difference in cellular hydrophobicity was observed (**Fig. 3E**). Notably, a progressive decrease in *Campylobacter* membrane potential was previously observed during bacterial incubation in sterile water and entering into VBNC state (Tholozan, Cappelier et al. 1999). Additionally, the ethidium bromide (EtBr) diffusion assay showed significantly increased membrane permeability in stressed cells compared to non-stressed cells (p <0.0001, **Fig. 3F**). Increased EtBr influx is a hallmark of stress and typically indicates that the cell is losing its ability to maintain membrane integrity or active efflux. Therefore, we next speculate that compromised membrane integrity will increase the production of bacterial vesicles.

### 3.2 Combined stress mediates the release of bEVs by *C. jejuni* Bf

Evidence has accumulated that bacterial vesicle production is influenced by both the genetic background of the strain and its growth conditions (Macdonald and Kuehn 2013, Toyofuku, Nomura et al. 2019, Kim, Seo et al. 2020, McMillan and Kuehn 2021, Malet-Villemagne and Vidic 2024). We isolated bEVs secreted by unstressed and stressed *C. jejuni* cells by ultracentrifuging the culture supernatants followed by Iodixanol-based density gradient centrifugation to remove protein aggregates, non-membrane proteins, and membrane debris, as shown in **Fig. S1**. Consecutive fractions were examined by TEM, which showed that bEVs were distributed in fractions containing 10-26% Iodixanol (**Figs. S1A** and **S1B**). bEVs isolated from *C. jejuni* appeared spherical with a clearly visible membrane composed of a lipid bilayer (**Fig. 4A**). The hydrodynamic diameter (*R*_*H*_) of bEVs measured by DLS showed that the density gradient purification step separated bEVs with *R*_*H*_ higher than 80 nm from a population of smaller particles (*R*_H_ ~20 nm), which most likely corresponded to secreted proteins and their aggregates (**Fig. S1C**). Nanoflow cytometry of purified bEVs stained with MemGlow membrane dye confirmed the purity of the bEVs with 84.8% intact vesicles and 15.2% opened or collapsed ones (**Fig. S1D**).

**Figure 4.**
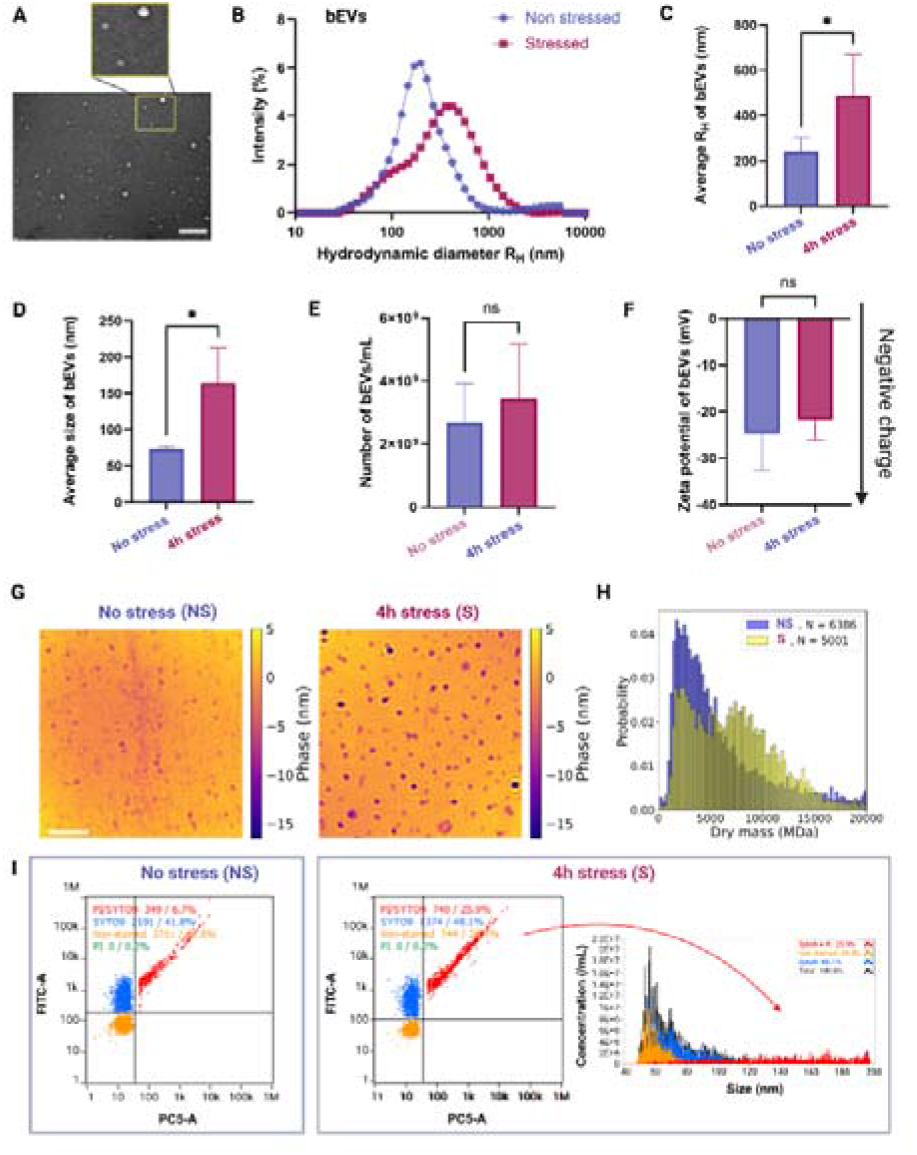
Under stress exposure, *C. jejuni* Bf cells produce larger and heavier extracellular vesicles carrying extracellular DNA. **(A)** Negative staining TEM micrographs of bEVs isolated from *C. jejuni* Bf culture. bEVs highlighted in the yellow square are shown enlarged above. The scale bar, 250 nm. **(B)** Hydrodynamic radius distribution of bEVs from non-stressed and stressed *C. jejuni* measured by DLS. **(C, D)** Average hydrodynamic radius and size of bEVs from the two groups obtained using DLS and NanoFCM, respectively. **(E)** Concentration of bEVs purified from a 60 mL culture of stressed and non-stressed *C. jejuni*, as determined by NTA. **(F)** Zeta-potential measurements reflecting the surface charge of bEVs secreted from stressed and unstressed *C. jejuni* Bf cells. **(G)** Quantitative phase images of bEVs from non-stressed and stressed *C. jejuni* imaged with Z-WIM. Scale bar, 5µm. **(H)** Dry mass distribution of bEVs from the two groups measured using phase images. **(I)** DNA content of bEVs estimated using the Syto9/propidium iodide (PI) assay showing the presence of external (PI) or internal DNA (Syto9). The right panel illustrates that large vesicles from stressed cells carried extracellular DNA. Data are presented as mean ± standard deviation from at least three independent experiments. Statistical significance was determined by Student’s t test. ns, not significant; * *p* < 0.05.

The size distribution of EVs, determined by DLS, showed that unstressed Bf cells produced a uniform population of vesicles, with a mean hydrodynamic diameter, R_H_, of 197.6 nm while stressed bacteria released a heterogeneous population comprising two main subpopulations with median R_H1_ of 92 nm, and R_H2_ of 361 nm (**Figs. 4B** and **S1B**). Due to the generation of larger vesicles under combined stress conditions, the mean bEV size determined by DLS and Nanoflow cytometry (nFCM) was higher for stressed (486.78 nm in DLS, 164.34 nm in nFCM) than for non-stressed bacteria (240.71 nm in DLS, 73.41 nm in nFCM) (**Figs. 4C** and **4D**). Together, these results indicate that stressed *C. jejuni* secrete bEVs that are approximately twice as large as those produced under microaerophilic conditions.

Across conditions, the mean vesicle yield in terms of particles per milliliter was similar between non-stressed and stressed samples with 2.6×10^9^ ± 1.25×10^9^ and 3.5×10^9^ ± 1.69×10^9^ particles/mL respectively, as determined by NTA (**Fig. 4E**). Despite their different size distributions, the surface charge of bEVs from non-stressed and stressed bacteria was comparable, with average values of −24.6 ± 8.0 mV and −21.7 ± 4.3 mV, respectively, as measured by zeta potential (**Fig. 4F**). Notably, compared to the bacterial cells themselves (mean surface charge of −10.5 mV; **Fig. 3D**), the bEVs exhibited a markedly more negative surface charge (mean of −23 mV). This difference may result from a selective enrichment of anionic membrane components during vesicle formation.

To further verify whether, under combined stress, larger bEVs were produced to transport more biological molecules compared to the control, we estimated their dry mass using a Quantitative Phase Imaging (QPI) technique called Z-WIM. The increased sensitivity of the Z-WIM method allows precise weighing dry mass of individual vesicles by their phase (Gentner, Rogez et al. 2024). As shown in the phase images, Z-WIM confirmed that stressed bacterial cells produced larger vesicles compared to unstressed cells (**Fig. 4G**). In addition, the larger bEVs were also heavier than those secreted by non-stressed cells, reflecting differences in their cargo (**Fig. 4H**). The dry mass of vesicles produced by non-stressed and stressed bacteria averaged 6063 MDa and 7141 MDa, respectively (**Fig. 4H**). For comparison, the dry mass of empty DOPC liposomes with a 100 nm diameter is around 120 MDa (Gentner, Rogez et al. 2024). Together, these results show that after consecutive thermal and oxidative stress, *C. jejuni* produces bEVs in similar number and with similar surface charge as non-stressed cells, but with increased sizes, R_H_, and dry mass, potentially reflecting differences in their cargo.

We previously showed that *C. jejuni* under microaerophilic conditions secretes a heterogeneous population of bEVs with about 42% carrying encapsulated DNA (Calzuola, Malet-Villemagne et al. 2025). Additionally, bacterial DNA and bEVs have been reported in human blood circulation (Hendrix and De Wever 2022). To determine whether stressed cells also use bEVs to export DNA, we labeled extracellular vesicles with Syto9 and Propidium Iodide (PI) and performed nFCM to measure their DNA content. Because PI cannot permeate intact membranes while Syto9 can, this dual staining allows discrimination between internal and external DNA. As expected, 41.8% of bEVs produced by non-stressed bacteria were stained only with Syto9, and 6.7% were double-stained with Syto9 and PI (**Fig. 4I**, left panel). In contrast, the population of bEVs secreted by stressed *C. jejuni* contained 48.1% Syto9-positive vesicles and 25.9% double-stained vesicles (**Fig. 4I**, right panel). No vesicles were stained only with PI, indicating that none displayed surface-associated DNA without also containing internal DNA. Moreover, DNA localization appeared to correlate with vesicle size under stressed conditions. DNA was mainly detected inside vesicles of median diameter (Syto9-stained, blue population), whereas larger vesicles (≥⍰80⍰nm) exhibited both internal and surface-associated DNA (Syto9/PI-stained, red population). In contrast, smaller bEVs (≤⍰60⍰nm) contained no detectable DNA (unstained, orange population in **Fig. 4I**), and they likely correspond to lipid-protein complexes. The presence of DNA in bEVs of Gram-negative bacteria is usually linked to explosive cell lysis. However, the secretion of larger vesicles containing chromosomal DNA is also associated with various abiotic stresses and/or biofilm formation (Toyofuku, Nomura et al. 2019).

### 3.3 *C. jejuni* produces bEVs with a distinct lipid signature under stress conditions

To further investigate the role of membrane remodeling in *C. jejuni* stress adaptation, we performed exploratory untargeted lipidomic analysis by mass spectrometry on both bacterial cells and their vesicles using a previously optimized chromatographic method for total lipid analysis (Moulin, Solgadi et al. 2015). We identified a total of 43 lipid species in both samples. All identified lipid species belonged to only 3 lipid classes: phosphatidylethanolamines (PE) (19 species), followed by phosphatidylglycerols (PG) (18 species) and lysophosphatidylethanolamines (LPE) (6 species). Surprisingly, the non-bilayer anionic cardiolipin species which are involved in respiration and osmotic stress responses were not detected in *C. jejuni* Bf, although cardiolipin is abundant in most bacteria (Arias-Cartin, Grimaldi et al. 2012). For each phospholipid species, the absolute peak intensity was compared between non-stressed and stressed conditions for both bacterial cells and bEVs (**Supplementary Table 1**). Peak intensities of PG, PE, and LPE phospholipids are represented as heatmaps in **Fig. 5A, B, and C**, respectively. To facilitate comparison between conditions, fold changes were calculated, and a fold change > 4 was considered significant.

**Figure 5.**
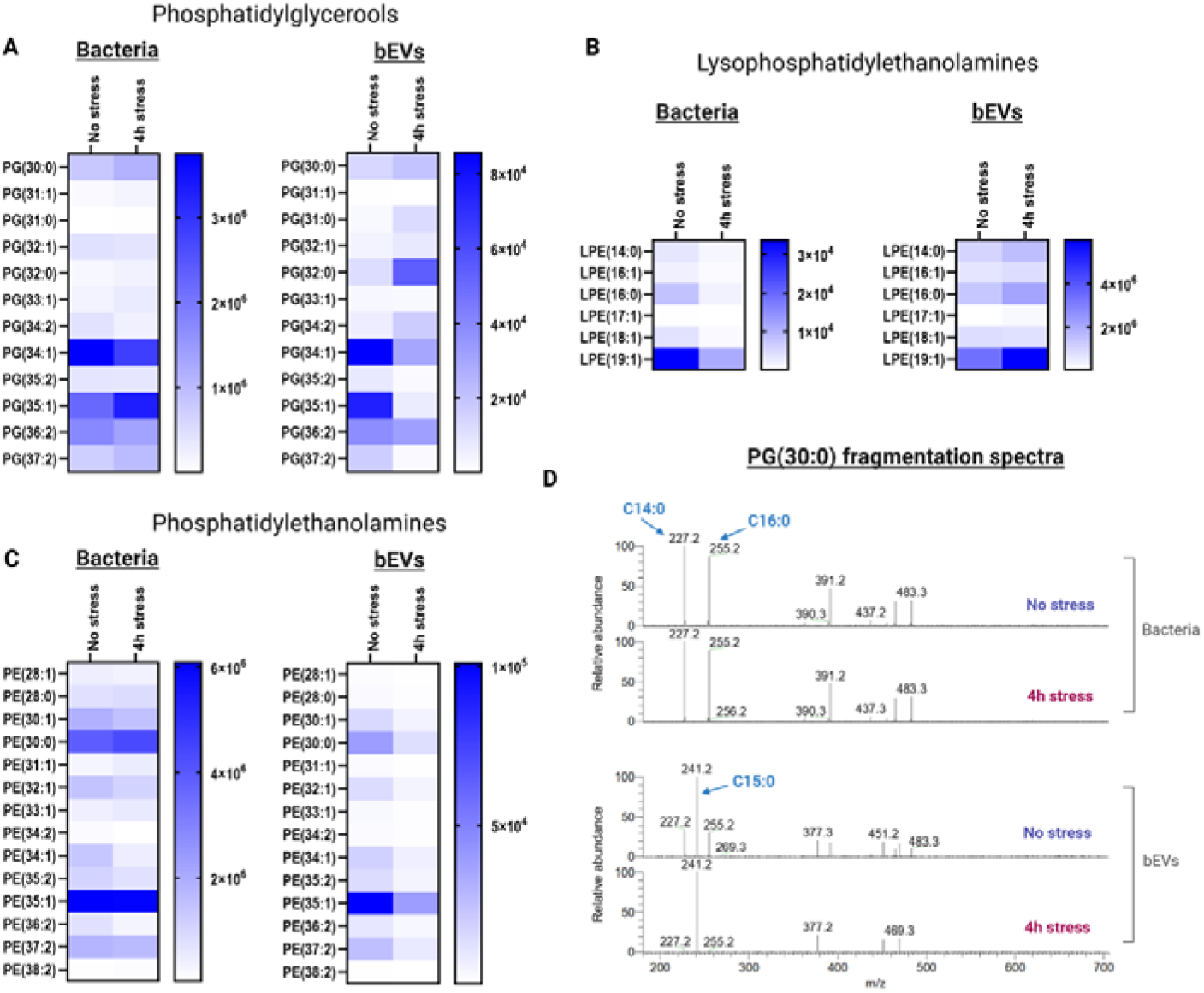
Stress exposure influences the lipid composition of cells and vesicles secreted by *C. jejuni* Bf. Phospholipids (PL) were extracted from the bacterial pellets or purified bEVs and separated by polarity. Individual species were identified in negative electrospray ionization mode (ESI-) using high-resolution mass detection. **(A, B, and C)** Heatmaps showing the phosphatidylglycerol (PG), lysophosphatidylethanoamines (LPE), and phosphatidylethanolamine (PE) phospholipids identified in all samples, and the relative abundance of the constituent fatty acids (ranging from 0% in white to 100% in dark blue). **(D)** Mass spectral profiles of the species PG(30:0) across all samples. The x-axis shows the m/z ratio of the ions, and the y-axis indicates their relative abundance. Note significant differences in PG profiles between the plasma membrane and bEVs.

In bacterial cells, no significant differences were observed in PG, PE, or LPE lipid species between stressed and non-stressed conditions (**Fig. 5A, B, and C**, left panels). In contrast, marked differences were detected in the lipid composition of bEVs derived from stressed versus non-stressed cells. Within the PG class, the PG(31:0) species (corresponding to PG(14:0–17:0) and PG(15:0–16:0) in the bEVs stressed sample) and the PG(32:0) species (corresponding mainly to PG(15:0–17:0) in both bEVs samples) were increased by 5-fold and 4.7-fold, respectively, in stressed compared to non-stressed bEVs (**Fig. 5A**, right panel). In addition, the only other PG species containing odd-chain fatty acids (C15:0 and C17:0), PG(30:0), is uniquely composed of PG(15:0-15:0) in stressed bEVs sample whereas it is composed of both PG(15:0-15:0) and PG(16:0-14:0) in non-stressed bEVs sample. Together, these results suggest that stress induces the production of membrane vesicles enriched in saturated odd-chain fatty acids in the PG lipid class by *C. jejuni*.

Interestingly, lipidomic analysis revealed that all PG phospholipids containing the C19:1 fatty acid were significantly reduced in bEVs membranes following stress. It should be noted that lipidomic analysis does not allow discrimination between unsaturated linear and cyclopropane lysophospholipids containing 19 carbons. But according to our fatty acid analysis (next section), this C19:1 fatty acid likely corresponds to cyclopropane C19. For several lipid species, this decrease was substantial in bEVs following stress, including an 11-fold reduction in PG(35:1) (corresponding to PG(16:0–19:1)) and an 8-fold reduction in PG(37:2) (corresponding to PG(18:1–19:1)), as shown in **Supplementary Table 1** and **Fig. 5A**.

Regarding lysophosphatidylethanolamines, the abundance of LPE(16:0) and LPE(18:1) was reduced after stress in bEVs membranes, with 4.7-fold and 4.9-fold decreases, respectively (**Fig. 5B**). Other LPE species (LPE(14:0), LPE(16:1), LPE(17:1), and LPE(19:1) were present at comparable levels in bEVs under the two conditions. Finally, no significant differences were observed within the PE lipid class between stressed and non-stressed conditions (**Fig. 5C**, right panel).

The most striking difference was observed in PG composition between bacterial cells and vesicles under both non-stressed and stressed conditions. Identical PG MS1 m/z displayed distinct fatty acid compositions in the cellular and vesicular membranes, with bEVs being enriched in PG species containing saturated odd-chain fatty acids, such as C15:0 and C17:0, which were not detected in *C. jejuni* cell membranes. As illustrated in **Fig. 5D**, PG(30:0) was only composed of PG(14:0–16:0) in the bacterial membrane, whereas it mainly consisted of PG(15:0–15:0) in vesicles (**Fig. 5B**). Similarly, PG(32:0) consisted largely of PG(16:0–16:0) in the stressed bacterial membrane but was mainly composed of PG(15:0–17:0) in vesicles. These findings indicate a specific remodeling of PG species during vesicle biogenesis.

Taken together, these results reveal a highly specific lipid composition of bEVs compared to bacterial membranes and show that stress triggers the production of bEVs with a distinct fatty acid signature in *C. jejuni*, underscoring vesicle biogenesis as an integral component of the bacterial stress response.

### 3.4 Combined stress drives unsaturated FA accumulation in *C. jejuni* and cyclopropane FA depletion in bEVs

We next quantified the fatty acids of *C. jejuni* Bf cellular and vesicle membranes under two environmental conditions. In both non-stressed and stressed cells, the bacterial cells contained seven main FAs: C14:0, C16:0, C16:1, C17:1, C18:0, C18:1, and C19 cyclo (**Fig. 6A**). This FA composition is consistent with the previously reported representative FA profile of most *C. jejuni* strains (Lambert, Patton et al. 1987), except for hydroxylated FAs such as 3-OH-C14:0, which was not detected. To further verify the absence of hydroxylated FAs in the *C. jejuni* Bf strain, we performed the extraction of FA methyl esters with diethyl/cyclohexane instead of diethyl/heptane (Dubois-Brissonnet, Trotier et al. 2016). However, this approach, which allows the extraction of hydroxylated FAs, produced FA profiles comparable to those obtained with the classical extraction (**Fig. S2**), further suggesting that the Bf strain does not produce hydroxylated FAs under the tested conditions. The fatty acid analysis showed that stressed cells had a significantly higher proportion of C17:1 compared to controls (2.76% vs. 0.68%, respectively; **Supplementary Table 1** and **Fig. 6B**), resulting in an overall increase in unsaturated FA content (**Fig. 6C**). No significant differences were detected in the levels of saturated or cyclopropane fatty acids.

**Figure 6.**
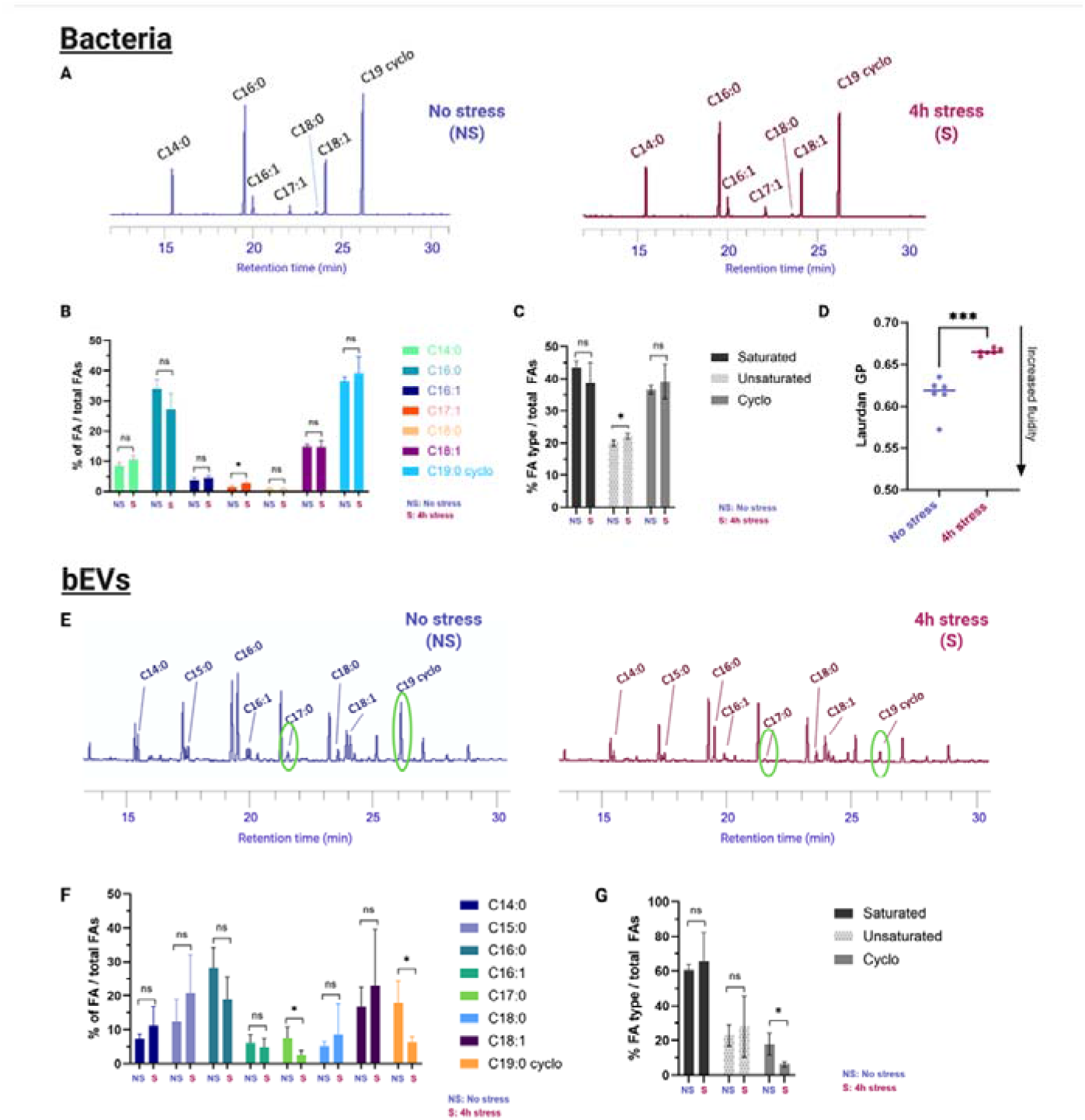
Fatty acid compositions of *C. jejuni* Bf cell and vesicle membranes change under stress exposure. **(A)** Gas chromatograms of fatty acids composing the membranes of stressed and non-stressed *C. jejuni* cells. **(B)** Relative abundance of detected fatty acids from the cell membrane, in %. **(C)** Relative abundance of fatty acid groups (saturated, unsaturated, cyclo-FA) in the membranes of stressed and non-stressed cells, in %. **(D)** Membrane fluidity measurements using Laurdan generalized polarization (GP) of stressed and non-stressed *C. jejuni* cells. Increasing relative Laurdan GP indicates membrane rigidification upon stress exposure. **(E)** Gas chromatograms of fatty acids composing the membranes of bEVs originating from stressed and non-stressed *C. jejuni* cells. **(F)** Relative abundance of all fatty acids composing the vesicle membrane, in %. **(G)** Relative abundance of fatty acid groups (saturated, unsaturated, cyclo-FA) in the membranes of vesicles released by stressed and non-stressed cells, in %. Peaks corresponding to fatty acids significantly less abundant in stressed vesicles are circled in light violet. Student’s t test was used to calculate p-values. ns, not significant, * *p* < 0.05, ** *p* < 0.01, *** *p* < 0.001, **** *p* < 0.0001.

Modification of membrane FA composition may preprogram membrane fluidity to maintain its function. Therefore, we measured the fluidity of the *C. jejuni* Bf membrane before and after consecutive stresses by measuring Laurdan generalized polarization (GP). The Laurdan probe intercalates into the membrane bilayer and displays an emission wavelength shift depending on the amount of water molecules in the membrane (Sánchez, Tricerri et al. 2007). Accordingly, an increase in the proportion of unsaturated FAs in the membrane leads to increased fluidity since the presence of double bonds in *cis* configuration introduces kinks in the carbon chain. Surprisingly, under combined oxidative and thermal stresses, the membrane of *C. jejuni* cells was significantly more rigid than that of non-stressed cells (**Fig. 6D**). This suggests that the membrane of stressed Bf is enriched in unsaturated C17:1 that adopts a *trans* configuration, resulting in an ordered phase. The increased rigidity of the bacterial membrane is an adaptive stress response that enables bacteria to save energy and survive.

At the vesicle level, eight FA species were identified: C14:0, C15:0, C16:0, C16:1, C17:0, C18:0, C18:1, and C19 cyclo (Fig. **6E-6G**). Notably, C15:0 was detected in bEVs but not in the plasma membrane. Moreover, our data indicate that the FA profiles of bEVs differ significantly from those of their parental cells, with bEVs displaying many unidentified peaks, often just before the major fatty acid peak (**Fig. 6A** and **6E**). We hypothesize that these unidentified peaks are modified fatty acids, likely differing at the branching level. For example, just before C16:0, there could be a C16:0 with a different branching pattern at the end of the chain, causing it to appear slightly later. This hypothesis still needs to be confirmed by including standards for branched antigens from *Campylobacter* or a closely related bacterium.

The relative abundance of C17:0 and C19 cyclo was significantly lower in bEVs originating from stressed cells, compared to bEVs from non-stressed cells (**Fig. 6E-6G**). This finding, together with specific lipid profiles (**Fig. 5C**), indicates that consecutive thermal and oxidative stress accumulates unsaturated odd-chain fatty acid C17:0 in the *C. jejuni* cell membrane. However, these stressed cells produce bEVs with lower abundance of odd-chain fatty acids C17:0 and C19 cyclo. Considering that odd-chain FAs alter lipid packing differently than even-chain FAs, our findings suggest the importance of C17:0 in fine-tuning membrane fluidity and helping *C. jejuni* to survive under temperature changes and oxygen level variations.

### 3.5 Stress impacts protein content in *C. jejuni* cells and their bEVs

We next investigated the proteomic characteristics associated with the phenotypic features of stressed *C. jejuni* Bf cells and their bEVs. Comparative nano-LC-MS/MS label-free analysis, after annotation with the UniProt database, identified a total of 660 proteins in control cells and 424 proteins in stressed cells (4 hours post-stress), respectively (**Supplementary Table 2**). The relatively low number of identified proteins is attributable to the limited reviewed annotation of the *Campylobacter* proteome (see section 2.12).

A principal component analysis (PCA) based on imputed LFQ (label-free quantification) intensities, requiring proteins to be present in at least 50% of the samples in one group showed that although there is a slight overlap between the two ellipses, an overall variance between the non-stressed and stressed samples is clearly observed (**Fig. 7A**). We may speculate that the small overlap reflects the non-simultaneous cell morphological changes observed in **Figs. 2A** and **2B**. Hierarchical clustering (HC) analysis revealed significant variations in protein profiles between conditions but not substantially between sample replicates (**Fig. S3**). The Venn diagram of the identified proteins showed that non-stressed and stressed *C. jejuni* cells contained 239 and 3 unique proteins, respectively, and more than 63% (421 out of 663) of the identified proteins were present in both (**Fig. 7B, Supplementary Table 2**). Notably, this analysis indicates that 36% proteins (239 out of 663) identified in the control cells were not found in stressed cells, strongly suggesting a decrease in bacterial metabolic activity. Among the uniquely expressed proteins in stressed cells, we found NAD^+^-dependent protein deacylase, which removes negatively charged acyl groups from lysine residues on proteins in *Campylobacter* (Jeter and Escalante-Semerena 2021). The production of this enzyme may contribute to the reduced bacterial negative charge observed in zeta-potential experiments (**Fig. 3D**).

**Figure 7.**
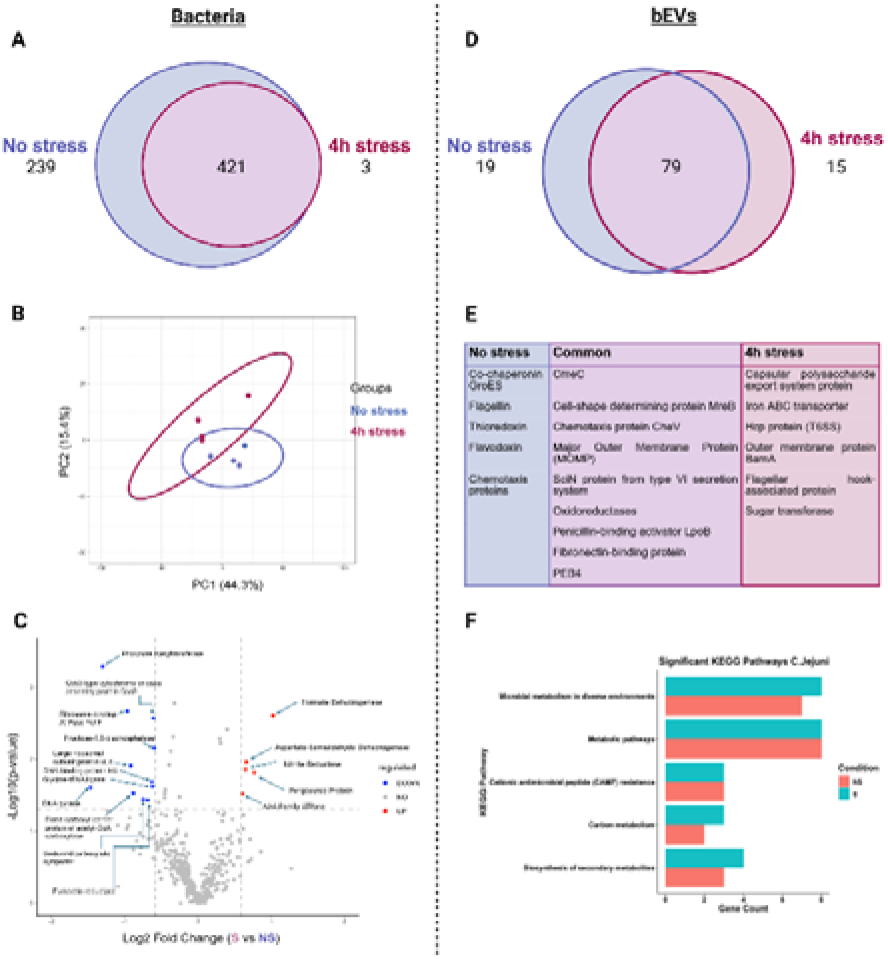
Stress exposure leads to protein content modifications in *C. jejuni* Bf. **(A)** Venn diagram showing the number of proteins identified in stressed and non-stressed *C. jejuni* cells. **(B)** PCA plot based on scaled LFQ intensities for identified proteins in stressed and non-stressed cells. Ellipses indicate the 95% confidence interval for each group. **(C)** Volcano plot illustrating differentially expressed proteins in stressed cells compared to controls (non-stressed). The x-axis shows the log2 fold change, and the y-axis shows −log10 (*p*-value). The significance threshold is set at adjusted *p*-value<0.05. **(D)** Venn diagram showing the number of proteins identified in bEVs from both stressed and non-stressed *C. jejuni*. **(E)** Table listing the major proteins found only in one bEVs group (stressed, or non-stressed) or in both bEVs groups. **(F)** Bar plot showing significant KEGG pathways, with a *p*-value threshold of 0.05. Data were obtained from independent experiments (biological replicates n = 5 for bEVs, n = 4 for cells).

In addition to the differentially detected proteins, 5 proteins were enriched, and 11 proteins were downregulated in the stress category compared to controls (using a p-value threshold of <0.05 and a fold change [FC] threshold of >1.5 or < 0.67) (**Fig. 7C**). Upregulated expression of proteins involved in energy metabolism: formate dehydrogenase (which catalyzes electron transfer via menaquinone and contributes to ATP synthesis) (van der Stel, Boogerd et al. 2017), and nitrate reductase (which transfers electrons from menaquinone to nitrite, supporting ATP synthesis) (Stahl, Butcher et al. 2012) may explain the high levels of ATP present in stressed cells (**Fig. 3C**). Aspartate-semialdehyde dehydrogenase, periplasmic protein, and AAA family ATPase were also upregulated. These proteins are linked to cell growth and metabolism, confirming that at the early stage of *C. jejuni* adaptation to oxidative and thermal stress, bacterial cells still perform biosynthesis and protein remodeling. Periplasmic proteins, in particular, have been shown to play a critical role in supporting *Campylobacter* survival in environmental reservoirs, including water and food matrices (Kemper and Hensel 2023). Interestingly, proteins of central metabolic processes, that are downregulated in stressed cells, such as phosphate acetyltransferase (TCA cycle), ribosome-binding ATPase YchF and glycine-tRNA ligase (protein synthesis), and biotin carboxyl carrier protein (fatty acid biosynthesis) (Birk, Wik et al. 2012, Hofreuter 2014, Landwehr, Milanov et al. 2021) are related to entry into the VBNC state (Zhao, Wang et al. 2016). Finally, the downregulation of key enzymes of anaerobic respiration: cbb3-type cytochrome C, which has high oxygen-binding affinity (Garg, Taylor et al. 2021) and fumarate reductase of the succinate:quinone oxidoreductase family (Weingarten, Taveirne et al. 2009), reflects the adaptation of the bacterium to oxidative stress associated with the general metabolic shift in dormant states.

Protein profiling of bEVs secreted by stressed (4 h post-stress) and non-stressed *C. jejuni* cells identified a total of 113 protein groups in five biological replicates for each condition (**Supplementary Table 2**). Protein distribution was balanced among the samples, with 70% (79 out of 113) of protein groups shared between both conditions, and 19 and 15 unique proteins found in vesicles from unstressed and stressed cells, respectively (**Fig. 7D**). *C. jejuni* Bf does not have a gene encoding cytolethal distending toxin, but many proteins identified in bEVs have pathogenic potential. The *Campylobacter* vesicle marker, MOMP (Malet-Villemagne and Vidic 2024, Calzuola, Malet-Villemagne et al. 2025), was detected in all tested samples (**Fig. 7E**). Notably, most unique proteins in bEVs of stressed cells are virulence factors located in the membrane and involved in transmembrane transport, including the iron ABC transporter (Palyada, Threadgill et al. 2004), capsular polysaccharide export system proteins (Riegert and Raushel 2021), flagellar hook-associated protein (Young, Davis et al. 2007), BamA, and Hcp (Malet-Villemagne and Vidic 2024). In contrast, unique proteins in bEVs from control cells are involved in central metabolism and protein folding, such as the oxidoreductase group (Banaś, Bocian-Ostrzycka et al. 2021), thioredoxin, GroES (Young, Davis et al. 2007) and chemotaxis proteins (Klappenbach, Negretti et al. 2021), as summarized in **Fig. 7E**. Despite the small number of annotated proteins, KEGG pathway (Kyoto Encyclopedia of Genes and Genomes) analysis confirmed cell metabolic adaptations to stress, with differentially expressed proteins involved in metabolic and biosynthetic interactions (**Fig. 7F**).

Overall, our proteomic analysis indicated that general stress responses and metabolic changes occur as *C. jejuni* copes with consecutive thermal and oxidative stresses. In addition, consecutive stress alters bEV cargo. Therefore, to distinguish the virulent potential of stressed and non-stressed *C. jejuni* cells and their bEVs, we examined the cytotoxic activities of bacteria and vesicles on eukaryotic epithelial cells.

### 3.6 Combined stress enhances the cytotoxicity of both *C. jejuni* cells and bEVs toward Caco-2 cells

Considering the changes observed in stressed and non-stressed bacteria, we investigated whether abiotic stress could increase the virulence of *C. jejuni* towards Caco-2 cells, and whether extracellular vesicles are involved in this process by transporting virulence factors.

We first evaluated the cytotoxic effect of *C. jejuni* culture supernatant at 20% (containing diluted bEVs) and purified bEVs at MOI 20 on Caco-2 cells in plates using the MTT viability assay. In this assay, cellular reduction of the tetrazolium dye MTT indicates cell viability (Vidic, Richard et al. 2016). While supernatants showed no reduction in cell viability, bEVs purified from non-stressed and stressed *C. jejuni* killed approximately 38% and 40% of treated Caco-2 cells within 24 hours (**Fig. 8A**). The decrease was not statistically significant between vesicles from non-stressed and stressed bacterial cells.

**Figure 8.**
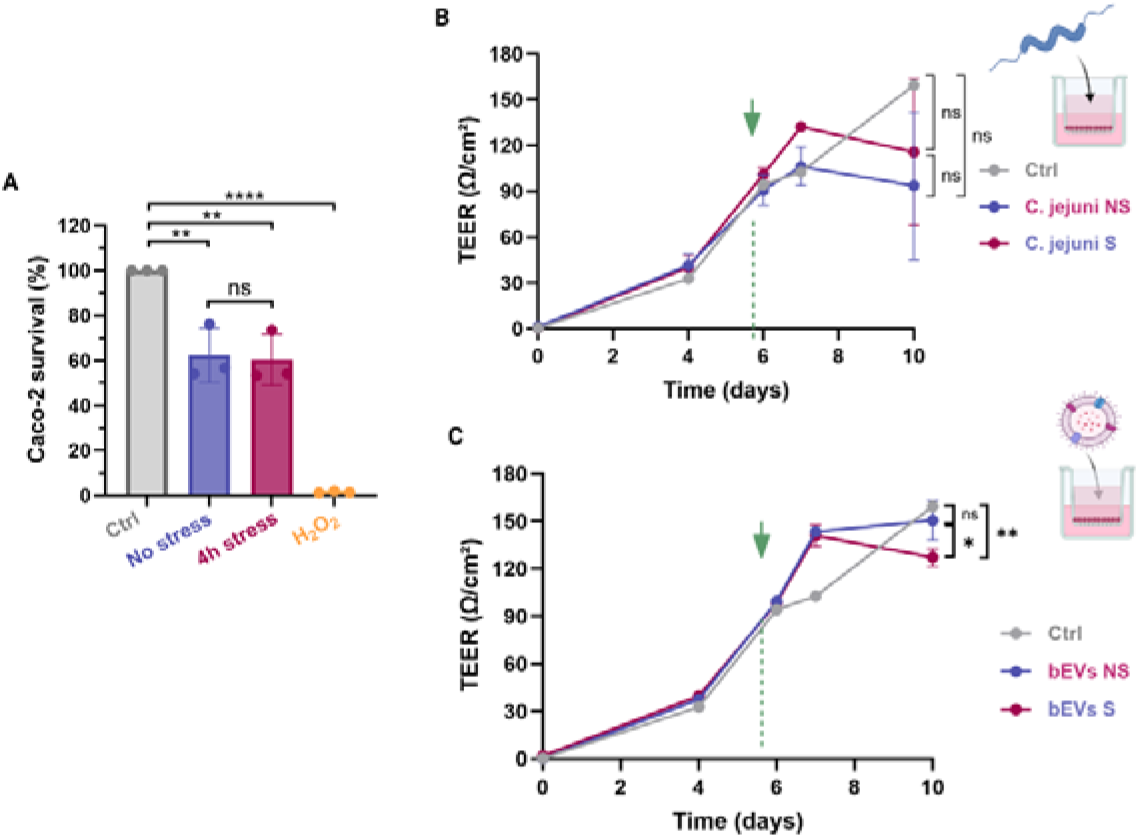
Exposure to stress increases the cytotoxicity of *C. jejuni* vesicles toward Caco-2 cells. **(A)** The toxicity of *C. jejuni* purified bEVs was evaluated by MTT test on a confluent monolayer of Caco-2 cells at a MOI 20. TEER (transepithelial electrical resistance) measurements of the Caco-2 cell monolayer were taken after challenge with either *C. jejuni* Bf cells **(B)** or purified bEVs **(C)** added to the apical channel. TEER experiments were performed on a day 6 when Caco-2 confluence was reached, as indicated by a green arrow with bEVs at a MOI of 20. Error bars indicate the standard deviation from three biological replicates. Student’s t test was used to calculate *p*-values. Data are presented as mean ± standard deviation from independent experiments (n=3). ns, not significant, * *p* < 0.05, ** *p* < 0.01.

We then evaluated Caco-2 tight junctions using a quantitative TEER assay, which assesses the epithelial barrier integrity (Calzuola, Malet-Villemagne et al. 2025). Caco-2 cells were cultured for 6 days to reach TEER values exceeding 100 Ω/cm^2^, indicating the formation of intact tight junctions and low paracellular permeability. Cell confluence was also confirmed by optical microscopy (**Fig. S4**). Stressed and non-stressed bacteria were then introduced into either the apical channel (**Fig. 8B**) or the basal channel (**Fig. S5**). In control wells, TEER values continued to increase until day 10, indicating ongoing cell growth through the end of the experiment. However, after inoculation in the basal channel of bacteria-treated wells, Caco-2 cells continued to grow for one day (until day 7), after which the TEER value began to decrease from 134.7 ± 8.3 Ω/cm^2^ to 86.3 ± 20.4 Ω/cm^2^ for non-stressed *C. jejuni* and from 148.9 ± 46.1 Ω/cm^2^ to 46.1 ± 32.4 Ω/cm^2^ for stressed *C. jejuni*, indicating that exposure to *C. jejuni* compromised the integrity of tight junctions in the Caco-2 cell monolayer. No significant difference was observed between the toxicity of stressed and non-stressed bacteria (p>0.05) (**Figs. S5** and **Fig. 8B** and **Supplementary Table 3**).

Next, we examined the effect of bEVs released by stressed and non-stressed *C. jejuni* on Caco-2 tight junctions by performing TEER experiments to evaluate their involvement in host infection. After bEV addition on day 6, cells continued to proliferate for one day. Subsequently, the TEER value in wells treated with stressed bEVs decreased from 141 ± 6.8 Ω/cm^2^ to 127 ± 5.5 Ω/cm^2^, whereas in wells treated with bEVs from non-stressed *C. jejuni*, the TEER values slightly increased from 141 ± 0.69 Ω/cm^2^ to 150 ± 12.5 Ω/cm^2^ (**Fig. 8C and Supplementary Table 3**). These findings suggest that vesicles from stressed *C. jejuni* cells disrupt more tight junctions in Caco-2 monolayers compared to bEVs from non-stressed cells which can facilitate bacterial infection of the host.

## 4. Discussion

In this study, we provide evidence that an aerotolerant *C. jejuni* Bf can survive and multiply under combined thermal and oxygen stress. During the early stage of adaptation (4 hours post-stress), the bacterial cells exhibited lipidomic, proteomic, and morphological responses, and released extracellular vesicles with increased toxicity to human epithelial barrier integrity.

*C. jejuni* is an enterohepatic species that inhabits the intestinal tract or gall bladder of its hosts. It commonly exists as a commensal bacterium in a wide range of animals, and its transmission to humans via food and water strongly depends on strain aerotolerance. Aerotolerance is one of the main strategies the bacterium employs to survive harsh stress conditions, along with biofilm formation and transition to the VBNC state (Murphy, Carroll et al. 2006, Oh, Kim et al. 2016). However, the aerotolerant state of Campylobacters remains a largely uncharacterized phenomenon. Studying combined stresses can predict *C. jejuni* behavior during poultry processing since cell history plays an important role in adaptation by induction of physiological responses involved in co-selection mechanisms (Huo, Xu et al. 2024, Léguillier, Pinamonti et al. 2024). *Campylobacter* strains that persist under harsh environmental conditions would provide a reservoir for human infections.

Multilocus sequence typing (MLST) of *C. jejuni* strains showed that aerotolerant strains were primarily classified into MLST clonal complexes CC-21 and CC-45, which are also the major complexes implicated in human gastroenteritis (Nielsen, Sheppard et al. 2010, Oh, McMullen et al. 2015). In contrast, *C. jejuni* strain Bf, studied here, belongs to the less common clonal complex ST-403, whose strains have been recovered from patients with gastroenteritis and Guillain–Barré syndrome (Islam 2010, Rodrigues, Pocheron et al. 2015, Gao, Tu et al. 2023). The Bf strain is known as an atypical aerotolerant strain and can be considered hyper-aerotolerant because it survives more than 24h exposed to ambient atmospheric conditions (**Fig. 3A**, and (Rodrigues, Pocheron et al. 2015)).

Following combined thermal and oxygen stress, the Bf strain showed a marked decrease in motility and remodeled its membrane to allow a shape change from S-shaped to coccoidal (**Fig. 2**; **Videos 1** and **2**). The strain is relatively robust, as not all cells adopted the coccoidal shape; some remained S-shaped and survived (**Fig. 2A**). Although it was previously shown that Bf can enter the VBNC state in its spiral form (Federighi, Tholozan et al. 1998), our results show that under combined stresses, Bf does not enter the VBNC state because its intracellular ATP level remains high and the cells continue to multiply slowly (Figs. **3A** and **3C**). While many stress conditions decrease ATP due to growth inhibition or metabolic slowdown, several studies indicate that transient or mild stress conditions can induce short-term increases or stabilization of ATP levels associated with adaptive metabolic responses and further behavior regarding subsequent stresses (Cohn, Ingmer et al. 2007, Dufour, Stahl et al. 2013, Kim, Chelliah et al. 2021). In line with this, our proteomics analysis indicated that proteins involved in energy metabolism, cell metabolism, and growth were upregulated following combined stress (**Fig. 7C**) enabling cells to multiply under harsh conditions.

We also showed that aerotolerance fine-tunes the physicochemical properties of the plasma membrane making it more rigid and potentially more mechanically resistant. Notably, the cell surface became less negatively charged (**Fig. 3D**), suggesting that the consecutive stresses modified expression of surface charged molecules such as lipooligosaccharide, proteins, and lipids. The S-to-coccoid transition in C. jejuni was previously shown to involve enzymatic remodeling and compaction of the peptidoglycan layer, but not extrusion or loss of the cell wall (Frirdich, Biboy et al. 2019). In this regard, we found that stressed cells show increased envelope permeability (**Fig. 3F**), which likely reflects modifications at the membrane level.

The major finding of this study regarding *C. jejuni* membrane adaptation to combined stresses is that, while maintaining the lipid composition of its plasma membrane, the bacterium can modulate the lipid composition of secreted bEVs, which depends greatly on environmental conditions. The greater negative charge observed for *C. jejuni* bEVs compared to the bacterial plasma membrane (**Fig. 4F** vs. **Fig. 3E**) may result from selective enrichment of anionic lipid components during vesicle formation. This is consistent with previous findings that membrane vesicles in Gram-negative bacteria often originate from regions enriched in negatively charged lipids, such as PG (Bonnington and Kuehn 2016, Roier, Zingl et al. 2016). These lipids preferentially accumulate in curved membrane domains, that favor vesicle budding. Functionally, a more negative surface charge could influence vesicle stability, interaction with divalent cations, or binding to host cell membranes, potentially modulating the biological activity of stress-derived bEVs.

The quantification of FAs in *C. jejuni* cells indicated that the odd-chain unsaturated C17:1 accumulated in the plasma membrane following stress where it likely adapted a *trans* conformation to increase envelop rigidity. The combined thermal and oxidative stresses may explain the paradoxical increase in membrane rigidity despite a higher proportion of unsaturated fatty acids (**Fig. 6C**). While heat stress generally induces bacteria to increase saturated or longer-chain lipids to counteract the fluidizing effect of temperature (Pedrotta and Witholt 1999, Heipieper, Meinhardt et al. 2003, Erimban and Daschakraborty 2022), oxidative stress can promote lipid peroxidation and alter the configuration of double bonds (e.g., conversion from *cis* to *trans*). *Trans* unsaturation packs more tightly than *cis* unsaturation, thereby reducing membrane fluidity. The interplay of these two stresses could, thus, lead to a lipid profile that maintains or even increases membrane rigidity, despite an apparent enrichment in unsaturated FAs. At the same time, the selective packaging of odd chain fatty acids C17:0 and C19 cyclo was limited in bEVs following stress, suggesting their importance in maintaining the appropriate membrane FA composition of the parental cells. The selective incorporation of FAs in bEVs was also observed in *Helicobacter pylori*, with an increase of C17:0 and a substantial reduction of C14:0, C18:1 and C19 cyclo in vesicles under antibiotic pressure (Krzyżek, Opalińska et al. 2025). Moreover, our findings indicate the contribution of lysophospholipids, which contain only one FA instead of two, in coping with stress. LPEs are virulence factors of *C. jejuni* and are required for bacterial motility in low-oxygen conditions (Cao, Brouwers et al. 2020, Cao, van de Lest et al. 2022). In particular, a short-chain LPE(14:0) was shown to be toxic to eukaryotic epithelial cells (Cao, van de Lest et al. 2022). Our lipidomic analysis revealed that LPE(14:0) was only slightly increased following stress in bEVs, but the abundance of LPE(16:0) and LPE(19:1) was highly reduced in bEVs of stressed cells compared to those of non-stressed cells (**Fig. 5C**). This maintaining LPEs in the bacterial plasma membrane has detrimental effects on membrane integrity, as previously shown for *Escherichia coli* survival under stress conditions (Rowlett, Mallampalli et al. 2017).

The ability of *C. jejuni* to cross polarized epithelial cells and invade them is considered a hallmark of *C. jejuni* pathogenesis and has been studied extensively (Russell and Blake Jr 1994, Monteville and Konkel 2002, Calzuola, Malet-Villemagne et al. 2025, Malet-Villemagne, Rizzotto et al. 2025). In the present study we found that both stressed and control *C. jejuni* cells can disrupt tight junctions and invade Caco-2 cells (**Fig. S4**). Nevertheless, the bEVs secreted following stress disrupted tight junctions in the Caco-2 monolayer more efficiently than control bEVs (**Fig. 8C**). The disruption of junctional complexes in epithelial cells, which form a barrier to prevent microbes from crossing into deeper tissues, is a key virulence strategy of *C. jejuni* and other diarrheagenic bacteria (Finlay and Falkow 1997, Malet-Villemagne and Vidic 2024). *C. jejuni* disrupts epithelial junctions mainly through HtrA-mediated cleavage of adherens junctions, opening the paracellular pathway for bacterial internalization (Malet-Villemagne and Vidic 2024). However, HtrA was not differentially expressed by stressed and non-stressed cells as determined by proteomic analysis to explain this finding (**Supplementary Table 2**). Our results suggest that invasion factors transported by vesicles, such as flagella hook-associated protein and BamA, detected only in bEVs from stressed cells (**Fig. 7E**), could facilitate bacterial penetration of the epithelium. This suggests that under thermal and oxidative stress conditions, *C. jejuni* releases vesicles to increase its ability to colonize and infect hosts. Therefore, based on our results obtained using the *C. jejuni* Bf strain, we propose that aerotolerant Campylobacters could employ a membrane fitness strategy to rapidly overcome stress conditions and increase their virulence.

## Supporting information

Supplementary Table 1 Lipidomics

Supplementary Table 2 Proteomics

Supplementary table 3 TEER data

## Supplementary material

The data supporting this article have been included as Source data, and Supplementary Information 1-3.

## Data Availability

The mass spectrometry proteomics data have been deposited to the ProteomeXchange Consortium via the PRIDE partner repository with the dataset identifier PXD074845.

## Acknowledgments

We acknowledge Mélissa Reyre and Suzana Krupova (Excilone, France) for access to the ZetaView instrument; Pascal Berto, Hadrien Robert and Anis Aggoun (Institut de la Vision, France) for expertise on ZWIM and DIPSI microscopy. We thank all the MicrobAdapt team for fruitful discussions, especially Philippe Gaudu, Clara Louche and Alexandra Gruss. This work has benefited from the facilities and expertise of MIMA2 facility (Microscopy and Imaging Facility for Microbes, Animals and Foods), INRAE Agroparistech, 78352 Jouy-en-Josas, France, https://doi.org/10.15454/1.5572348210007727E12.

## CRediT roles

Conceptualization: JV; Data curation: JMV; Formal analysis: JMV, BP, AS; Funding acquisition: JV, GT; Investigation: JMV, RD, KG, BP, CP, AS, VC, ZZ; Methodology: JMV, JV, RD, ZM, KG, BP, AS, FDB, CP, VC, MDP, ZZ, GT; Project administration: JV; Software: RD, ZM; Resources: JV, FDB, MDP; Supervision: JV, ZM, GT; Validation: JMV, JV; Visualization: JMV, JV; Writing – original draft : JMV, JV; Writing – editing and review : all.

