## Supplementary Information for "Versatility of *Campylobacter jejuni* Bf extracellular vesicles in regulating adaptation and virulence under combined thermal and oxidative stress"

Summary:

Figure S1: Isolation and purification protocol of EVs.

Figure S2: Fatty acid profiles of *C. jejuni* Bf cells identified by GC-MS.

Figure S3: Heat map of *C. jejuni* Bf proteins.

Figure S4: Optical microscopy images of Caco-2 monolayers on Transwell membranes before TEER measurement.

Figure S5: TEER (transepithelial electrical resistance) measurements of *C. jejuni* cells added on Caco-2 cells by the apical channel

**Fig. S1.** Isolation and purification protocol of EVs. After supernatant centrifugation, the pellet contains bEVs but also protein aggregates (A). To eliminate protein aggregates, the pellet is resuspended in PBS and added on a 3-layer (10%, 26%, 45%) discontinuous iodixanol gradient, shown in (B). Vesicles were found in fractions 2, 3, 4 and 5 (very few), as shown on the electronic microscopy pictures. Scale bars = 500 nm. Measurement of hydrodynamic diameter R_H_ of particles present in the supernatant before and after gradient by DLS confirm the elimination of protein contaminants from the vesicle sample (C). Finally, a MemGlow membrane staining of the particles analyzed in NanoFCM show a minimal purity of 85% (D).

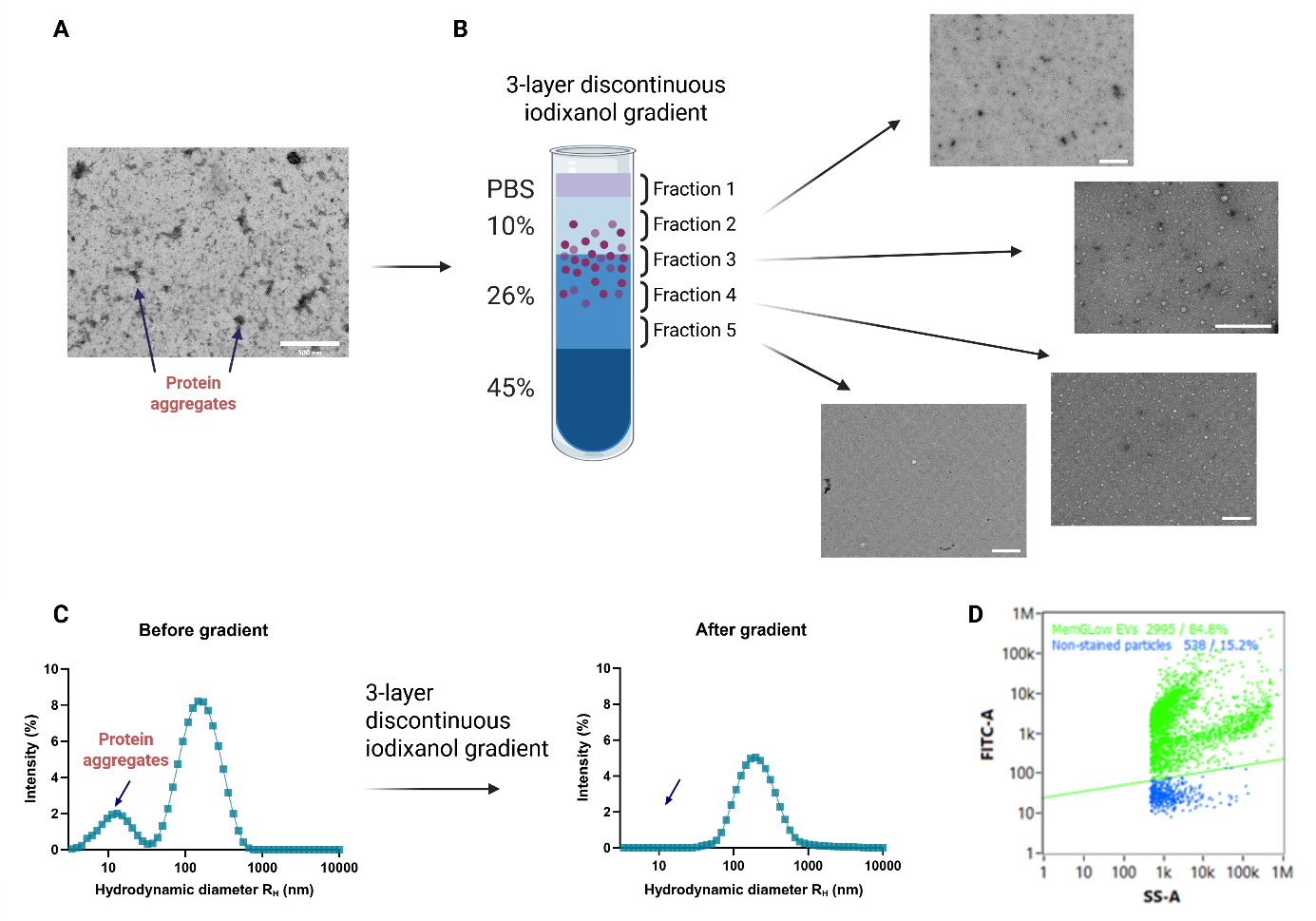

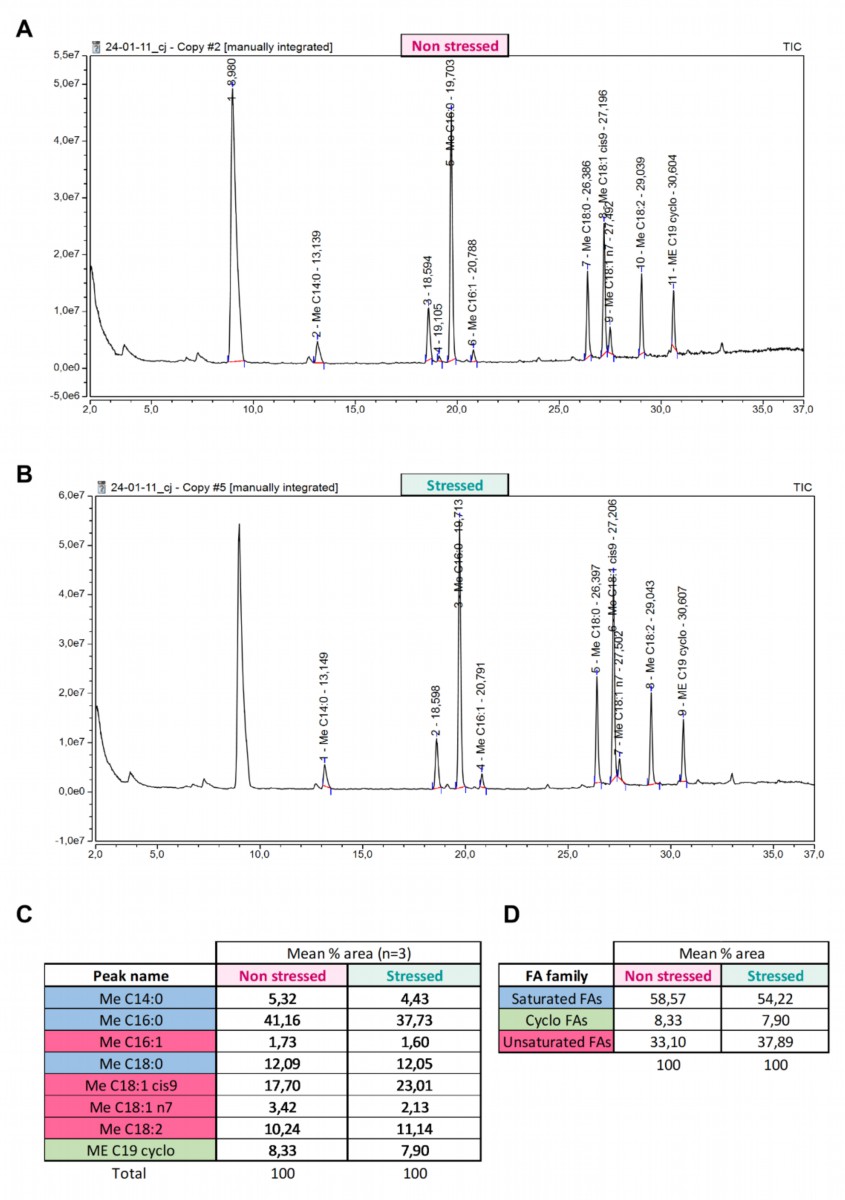

**Fig. S2.** Fatty acid profiles of *C. jejuni* Bf cells after extraction using diethyl/cyclohexane (allowing extraction of hydroxylated FAs) and identified by GC-MS. Fatty acids profile of non-stressed (A) and stressed (B) bacteria are presented. The relative abundance of each fatty acid was calculated using peak area (in % of total peak area). Mean relative abundance (n=3) of fatty acids (C) and fatty acid groups (D) in non-stressed and stressed cells.

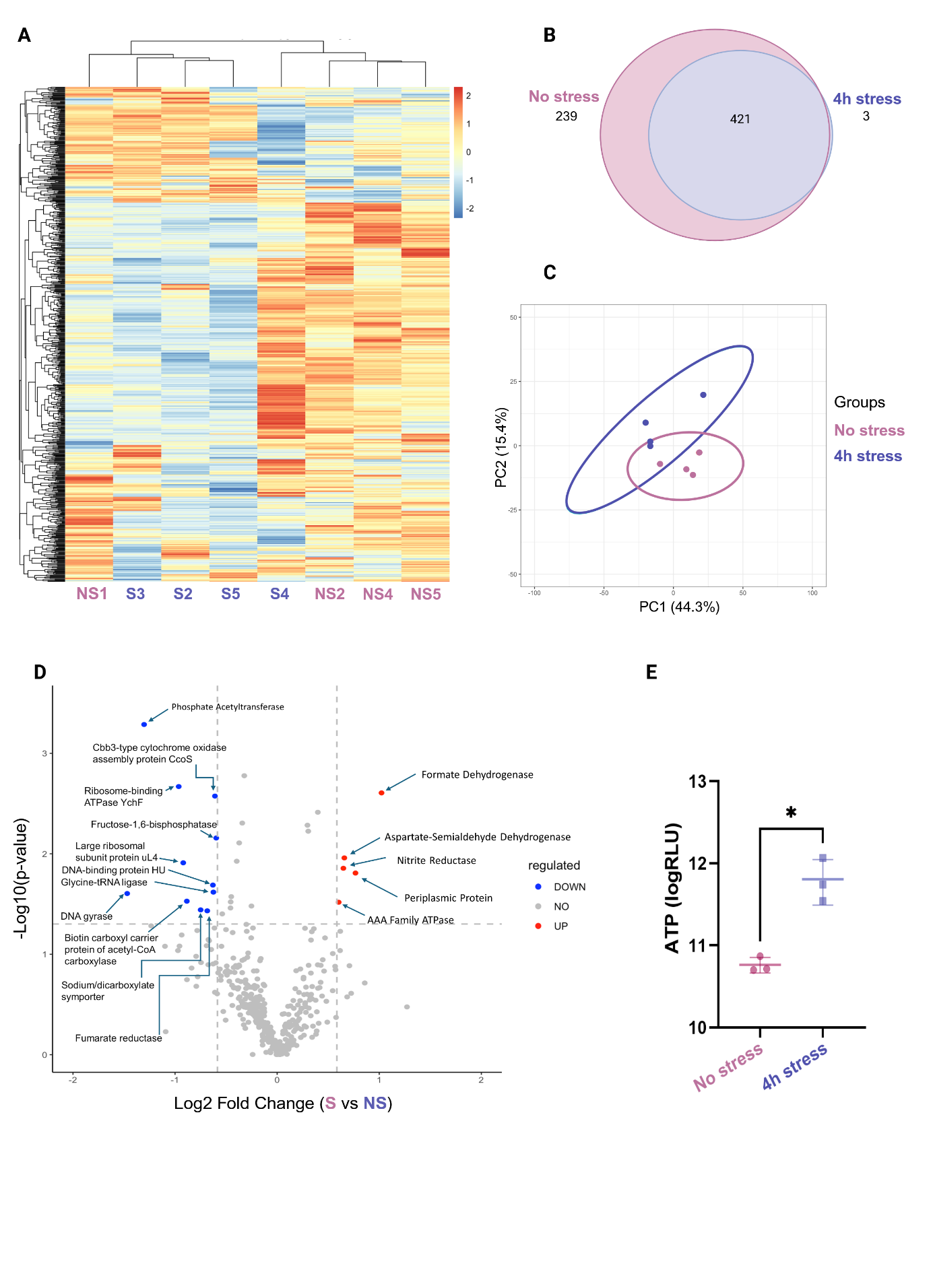

**Fig. S3.** Interactive cluster heatmap representing *C. jejuni* Bf proteins identified in non-stressed and stressed cells. 4 biological replicates were analyzed for each condition (n=4). Heat map was realized based on mean z-scored protein LFQ intensities of 1084 differentially expressed proteins (ANOVA FDR < 0.05) after unsupervised hierarchical clustering.

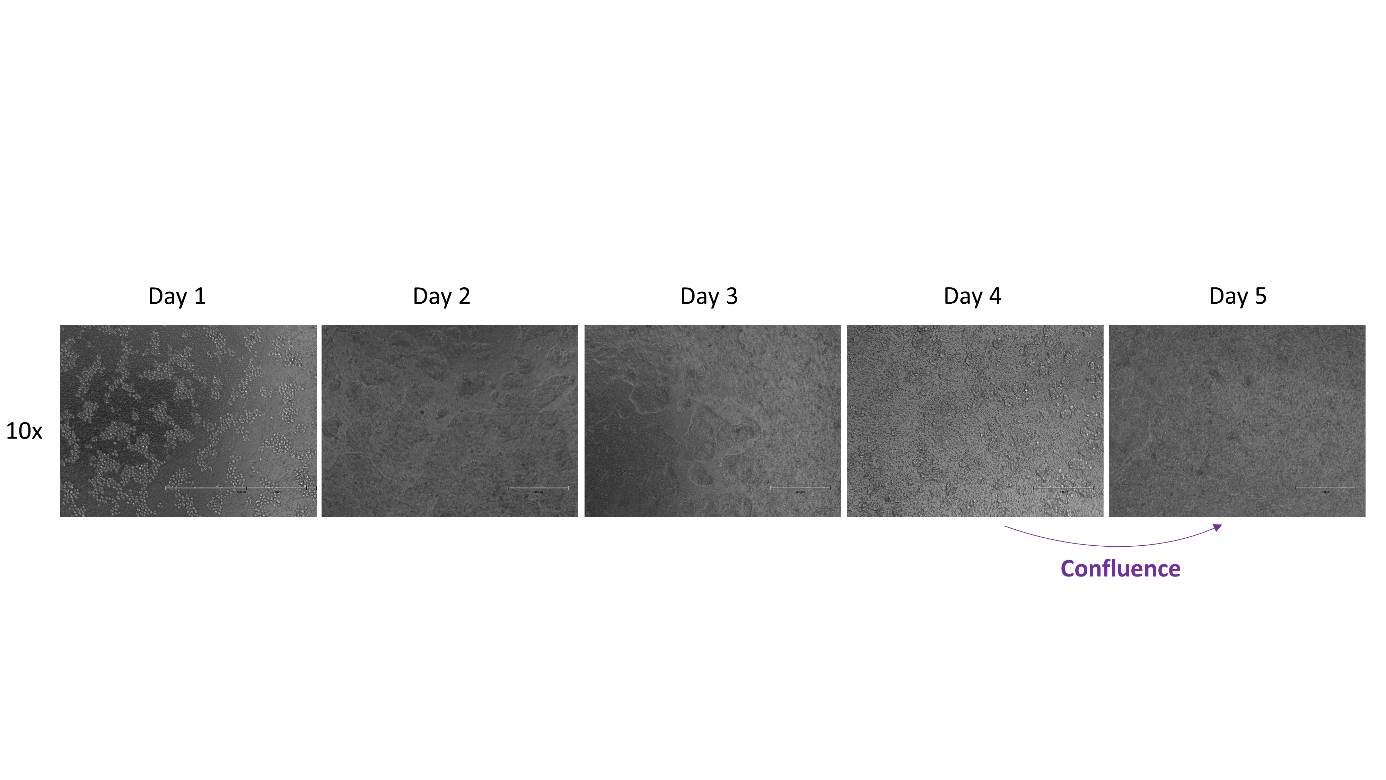

**Fig. S4.** Morphological evolution of Caco-2 monolayers on Transwell membranes before TEER measurement. Microscopic validation of cell confluence, obtained between day 4 and day 5. Brightfield images are presented with scale bars.

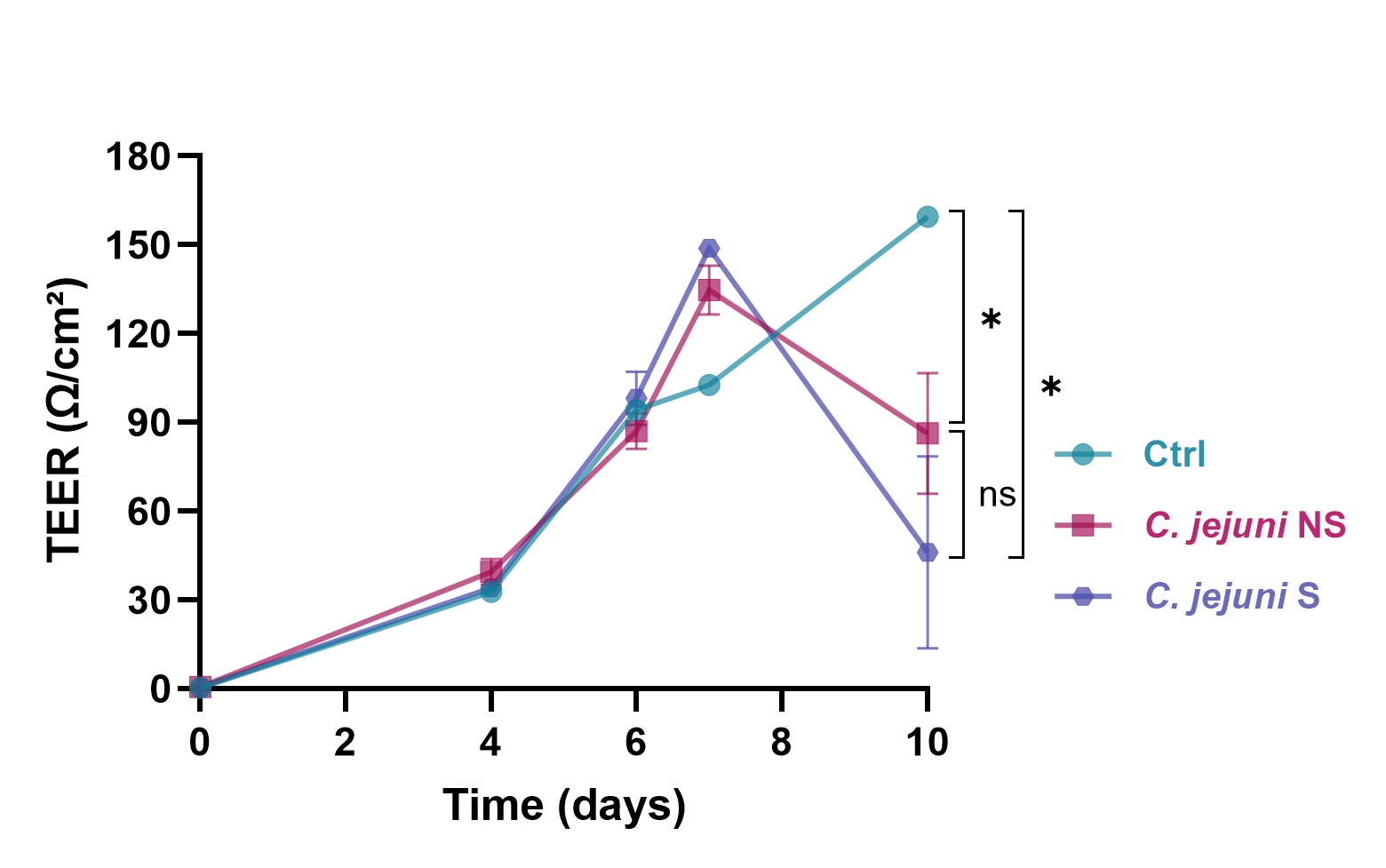

**Fig. S5.** TEER (transepithelial electrical resistance) measurements of *C. jejuni* non-stressed (NS) or stressed (S) cells added on Caco-2 cells by the basal channel. Error bars represent the SD from three independent experiments (n = 3). Confluence was reached at day 6, and is marked with a green arrow.

**Videos S1. and S2.**
